# A conserved function of corepressors is to nucleate assembly of the transcriptional preinitiation complex

**DOI:** 10.1101/2024.04.01.587599

**Authors:** Alexander R. Leydon, Benjamin Downing, Janet Solano Sanchez, Raphael Loll-Krippleber, Nathan M. Belliveau, Ricard A Rodriguez-Mias, Andrew Bauer, Isabella J. Watson, Lena Bae, Judit Villén, Grant W. Brown, Jennifer L. Nemhauser

**Affiliations:** Department of Biology, University of Washington, Seattle, 98195, USA; Department of Genome Sciences, University of Washington, Seattle, 98195, USA; Department of Biochemistry and Donnelly Centre, University of Toronto, Toronto, Ontario, CA

**Keywords:** Corepressors, Transcription initiation, TOPLESS (TPL), DSIF, SPT4/SPT5 complex, SPT6, TFIID, TAF5, proximity labeling, Reporter Synthetic Genetic Array, Integrases

## Abstract

The plant corepressor TPL is recruited to diverse chromatin contexts, yet its mechanism of repression remains unclear. Previously, we have leveraged the fact that TPL retains its function in a synthetic transcriptional circuit in the yeast model *Saccharomyces cerevisiae* to localize repressive function to two distinct domains. Here, we employed two unbiased whole genome approaches to map the physical and genetic interactions of TPL at a repressed locus. We identified SPT4, SPT5 and SPT6 as necessary for repression with the SPT4 subunit acting as a bridge connecting TPL to SPT5 and SPT6. We also discovered the association of multiple additional constituents of the transcriptional preinitiation complex at TPL-repressed promoters, specifically those involved in early transcription initiation events. These findings were validated in yeast and plants through multiple assays, including a novel method to analyze conditional loss of function of essential genes in plants. Our findings support a model where TPL nucleates preassembly of the transcription activation machinery to facilitate rapid onset of transcription once repression is relieved.

## Introduction

During development, efficient and coordinated switching between OFF and ON gene states is essential for cell fate determination and morphogenesis. This transcriptional reprogramming requires regulation of both activation and repression.

Failure of transcriptional repression leads to catastrophic defects in development such as complete loss of body plan organization^1,2^, oncogenesis and even death^3,4^. Despite this importance, the exact mechanism by which repressors enact and maintain repression, and especially their complex interactions with the highly conserved transcriptional activation machinery, remains elusive. Corepressors are one group of repressor proteins that are recruited by DNA-binding transcription factors and repress by recruiting negative regulators of transcription and/or inhibiting the active components of transcription such as RNA-Polymerase II (Pol-II). Transcriptional corepressors are found across all eukaryotes and in several structurally related families including the animal SMRT (silencing mediator of retinoic acid and thyroid hormone receptor) and NCoR (nuclear receptor corepressor) complexes^5,6^, the yeast Tup1^7–9^ and its homologs Drosophila Groucho (Gro) and mammalian transducing-like enhancer (TLE)^10^.

In land plant lineages, there has been an expansion of the Gro/TLE-type corepressor family, including TOPLESS (TPL), TOPLESS-RELATED (TPR1-4), LEUNIG (LUG) and its homolog (LUH), and High Expression of Osmotically responsive genes 15 (HOS15)^1,11–14^. All members of these families share a general structural homology, where the N-terminal domain contains a conserved protein dimerization domain known as the LIS1 homology (LisH) domain^15,16^. At the C-terminus of these proteins are WD40 repeats that form beta-propeller structures that are involved in many diverse protein-protein interactions^17,18^. TPL/TPR activity is essential for development, responses to the environment, and immunity^11,19^. In our previous work, we recapitulated the Auxin Response Circuit including TPL from the model plant *Arabidopsis thaliana* in *Saccharomyces cerevisiae* (yeast) (*At*ARC^Sc,20^). In the *At*ARC^Sc^, an auxin-responsive transcription factor (ARF) binds to a promoter driving a fluorescent reporter that can be quantified by flow cytometry (Figure S1A). In the absence of auxin, ARF activity is repressed by interaction with a full-length Auxin/Indole-3-Acetic Acid protein (hereafter IAA protein) fused to the N-terminal domain of TPL^20^. Notably, the N-terminal domain of TPL is sufficient for strong repression of transcription in yeast and in plants, can be subdivided into two distinct transcriptional repression domains: Helix 1 (H1) which is part of a LisH domain and Helix 8 (H8)^21^. H8 makes direct contact with Mediator 21 (Med21) and Mediator 10 (Med10), likely leading to assembly of the entire Mediator complex at a promoter while simultaneously inhibiting the transition to active transcription^21^. Despite its broad functional conservation^22^ and importance to TPL activity, the mechanism of H1/LisH-based repression on transcription is unknown.

Structure-function studies indicate that the modes of TPL action are distinct from other well-characterized modes of repression in plants and fungi, such as those that rely exclusively on epigenetic marks on histones or DNA methylation^23^. TPL appears to act as a priming agent for transcriptional activation, facilitating the assembly of Mediator and other components of the transcriptional preinitiation complex (PIC) at promoters bound by transcriptional activators^22,24^. This form of repression is reminiscent of RNA Polymerase II proximal promoter pausing in metazoans^25^. During transcriptional pausing, Poll II initiates transcription but then stalls as a result of a metazoan-specific protein called Negative Elongation Factor (NELF) interacting with the DRB Sensitivity Inducing Factor (DSIF) complex. The DSIF components SPT4 and SPT5 are conserved across eukaryotes^26^, and in concert with SPT6^27^, are crucial elongation factors that are associated with Pol-II at all points during transcriptional activation and elongation^28–31^.

While these factors are best known for their roles in elongation and pausing, they were originally identified in yeast as factors critical for promoter identification and transcriptional initiation^32,33^. Recent work has underscored the role of DSIF/SPT components specifically in transcription initiation in multiple organisms, such as SPT5 in metazoans^34^ and plants^35^, and SPT6 in yeast^36^ and plants^37^. In NELF-based pausing, and by extension in TPL-based priming, the transition from OFF to ON state is likely to be accelerated by the pre-assembly of necessary protein complexes.

In the current study, we leveraged the power of yeast genetics at genomic scale to uncover the molecular mechanism of TPL repression. We applied an unbiased proximity labeling approach in *At*ARC^Sc^ strains to map the physical interaction landscape of TPL in its repressed state at a single synthetic locus. In parallel, we performed genome-wide reporter synthetic genetic array (R-SGA) screens to find genes whose function was required for maintaining repression by H1 alone or the combination of H1 and H8 together. Among the most prominent candidates showing both physical and genetic TPL interactions were SPT4, SPT5, and SPT6. We further mapped the protein interaction interface between TPL and SPT4 in both yeast and plants through protein interaction assays, and found that SPT4 acts as a bridge to SPT5 and SPT6, both of which are required for H1-based repression. In addition, we validated TPL interactions with TAF5, a critical component of the PIC and showed that multiple PIC components are required for maintenance of a repressed state. Given the broad conservation of TPL interacting proteins across eukaryotes, we tested the function of TPL in human cells, and found that TPL was able to repress transcription when fused to dCas9. Finally, we developed a novel integrase-based method for conditional loss of function of essential genes in plants to test the impact of TPL interactors on an endemic TPL-repressed developmental process. Collectively, our findings support a model where TPL enables the stable preassembly of the PIC and associated initiation factors at promoters already associated with transcriptional activators. This mode of repression facilitates rapid onset of transcription once repression is relieved, as well as rapid re-establishment of the repressed state as the concentration or activity of inhibitory factors, such as Aux/IAAs, increases.

## Results

### Identifying the TPL protein interaction network in yeast through APEX2 proximity labeling

To understand how TPL represses transcription, we took an unbiased proximity labeling approach using the recently characterized ascorbate peroxidase 2 (APEX2) system for rapid labeling in *Saccharomyces cerevisiae*^38,39^. We generated a version of the *At*ARC*^Sc^* with the TPL N-terminal domain (amino acids 1-188; TPLN188) C-terminally fused to the APEX2 peroxidase^40^, to label proteins that interact with TPL using a small clickable alkyne-phenol probe (Alk-Ph, Figure 1C). As our previously described *At*ARC*^Sc^* was localized to chromatin by a protein fusion of TPL to IAA proteins^20^, we first demonstrated that robust transcriptional repression was possible using an unfused TPLN188 and IAA14. In this test, each protein was independently expressed and interacting via the Ethylene-responsive element binding factor-associated Amphiphilic Repression (EAR) motif on IAA14^11,41,42^. We observed that TPLN188-APEX2 performed equally well as a repressor as TPLN188 alone (Figure 1C,D). We performed our proximity labeling on yeast cultures as described in Li et al., 2020^39^ where the use of a small, cell-permeable and clickable Alk-Ph probe is the substrate for the APEX2 enzyme. Cells were subsequently lysed and biotin was attached to Alk-Ph labeled proteins through click chemistry. We observed robust and specific labeling of yeast protein extracts by the TPLN188-APEX2 fusion protein (Figure 1E,F). We observed weak but detectable background bands (Fig 1F, arrows) in the samples where H_2_O_2_ was omitted as a negative control, which likely correspond to endogenous peroxidase activities in yeast as was previously reported in Li et al., 2020^38,39^.

**Figure 1.**
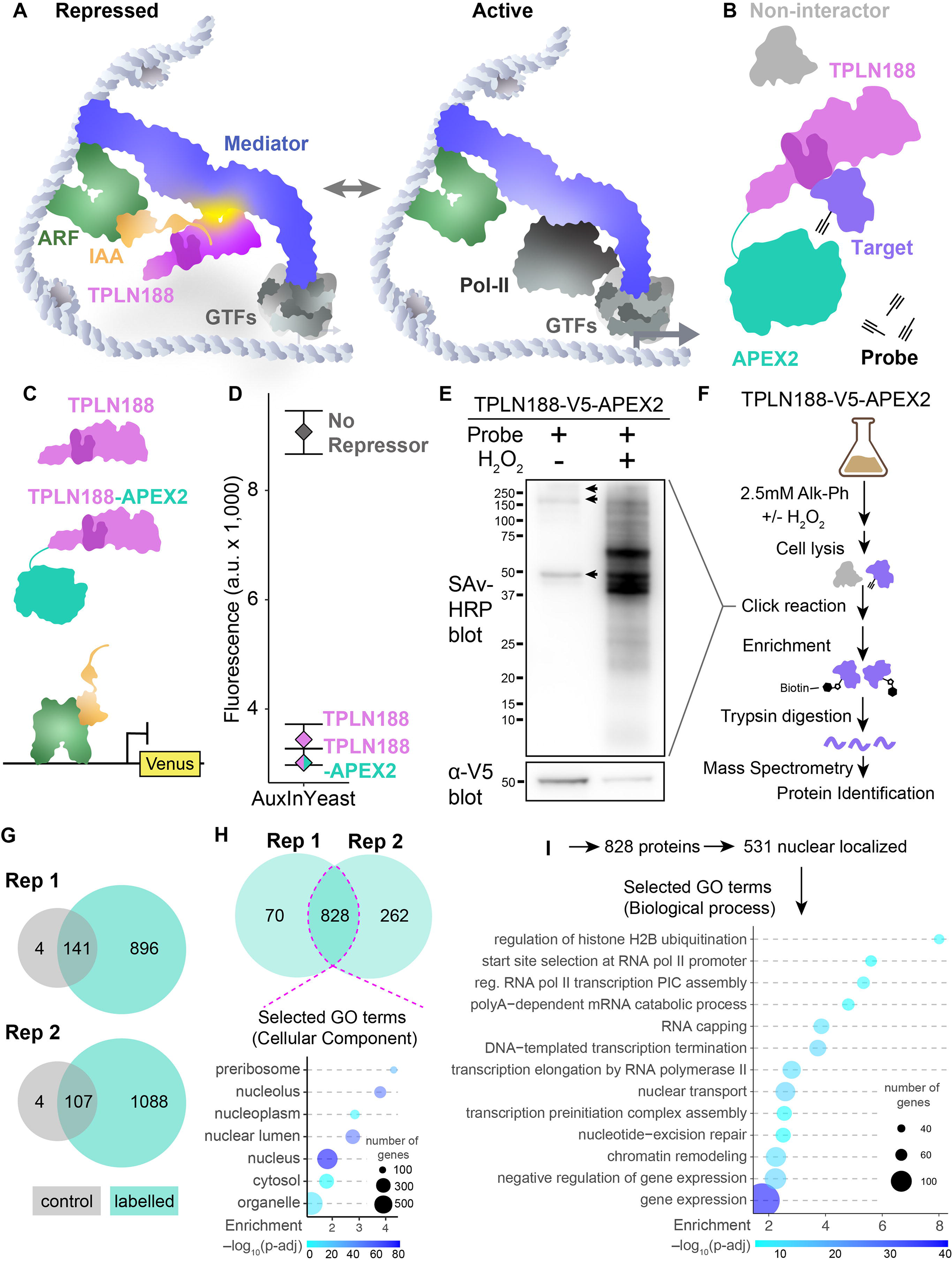
Identifying TPL proteins interactors in yeast through APEX2 proximity labelling. A. Schematic of the native auxin nuclear response components switching from repressed to active as we currently understand it. ARF – Auxin response factor (TF), IAA – AUX/IAA proteins, TPLN188 – the N-terminal domain of TPL, with the LisH highlighted in a darker shade, GTFs – general transcription factors, Mediator – the mediator complex, and Pol-II – RNA polymerase II. **B.** Schematic of the TPLN188- APEX2 fusion protein and its intended use in proximity labelling TPL-interacting proteins. **C.** Schematic of the use of TPLN188 (purple) or TPLN188-APEX2 (purple/teal fusion protein) in the AtARCSc in a design unfused to the IAA (orange). The ARF (green) is recruited to a promoter driving Venus (yellow) for detection by flow cytometry. **D.** Flow cytometry experiments of the TPL proteins indicated. Every point represents the average fluorescence of 5–10,000 individually measured yeast cells (a.u.: arbitrary units). **E.** Streptavidin-HRP blot analysis to compare TPLN188-APEX2-labeling efficiency in yeast with Alk-Ph with and without H_2_O_2_. Alkyne-modified proteins were ligated with azide-(PEG)_3_-biotin via click reaction. Molecular weight standards are shown in kDa. Arrows indicate endogenously biotinylated proteins. Bottom: 18α-V5 western blot showing the expression of TPLN188-APEX2. **F.** Workflow of protein-level proteomic experiments. Negative controls are yeast cells treated in the absence of the H_2_O_2_. Experiments were performed with two replicates for each condition. **G.** Venn diagrams showing the numbers of proteins identified across replicated proteomic experiments ± H_2_O_2_, gray – control, labelled – teal. **H.** Venn diagrams showing the numbers of proteins overlapping between the plus probe experiments that are not found within the control condition. Dotted pink line indicates the overlap between experiments which were taken for further analysis. Bottom: Selected Gene Ontology (GO) analysis for Cellular Component of all 828 proteins identified within the pink line. **I.** Selected gene ontology analysis for nuclear proteins identified within the pink line.

We purified biotinylated proteins by streptavidin affinity purification and prepared samples by Trypsin digestion for mass spectrometry (see methods). We performed two replicates of this protocol and observed significant overlap between replicates (Figure 1H, dotted magenta line). Several proteins served as internal positive controls in our experimental design: (1) the AUX/IAA protein which recruits TPL to chromatin, (2) the TPLN188-APEX2 fusion protein itself, and (3) the ARF transcription factor. All three were found to be highly enriched in our mass spectrometry identification, suggesting a successful and fairly specific activity that we can benchmark by relative enrichment compared to these controls (Figure S1B). The auxin receptor (AFB2) was identified with greater enrichment than ARF19 (Figure S1B), suggesting that, even in the absence of auxin, these proteins are localized in close proximity to the other components of the auxin response machinery.

As the TPLN188-APEX2 fusion protein is expressed ubiquitously from the yeast GPD promoter, proximity labeling should capture a snapshot of interacting proteins throughout the lifecycle of the protein, including any residence time in the cytoplasm after translation. However, we observed that a majority of identified proteins (64.1%) are localized to the nucleus (531/828, Figure 1H). Of these nuclear localized proteins, there was enrichment in GO terms connected to transcription (Figure 1I).

### Identifying the TPL genetic interaction network in yeast through Reporter Synthetic Genetic Array (R-SGA)

APEX proximity labeling of targets identified a large number of physical TPL- interactor proteins with an unknown relationship to repressor function. We set out to perform a high throughput test for the genetic interaction between endogenous yeast genes and the TPL N-terminal repressor domain using a Reporter Synthetic Genetic Array (R-SGA) approach^43^. We have previously engineered a version of the *At*ARC*^Sc^* that carries the entire auxin response circuit on a single CEN-type plasmid, which we named the Single Plasmid Auxin Response Circuit or SPARC^21^. In the SPARC, selected truncations of the TPL N-terminal domain are encoded as fusions to the IAA14 protein (Figure 2A, *TPLN-IAA14* - purple). Also encoded on the SPARC are the auxin receptor (Figure 2A, *AFB2* - blue), the LEU2 selectable prototrophy gene (Figure 2A, *LEU2* – gray), the auxin response factor transcriptional activator (Figure 2A, *ARF19* - green), and the Venus fluorescent reporter under the control of a promoter with a well-characterized ARF binding site (Figure 2A, *Venus* - yellow). Consolidation of parts into the SPARC allows its use in the R-SGA approach, as it can be used as a single selectable entity that acts as a reporter of repression, where modulation of repression of the Venus reporter in a mutant background will uncover genetic interactions between endogenous genes and TPL-based repression.

**Figure 2.**
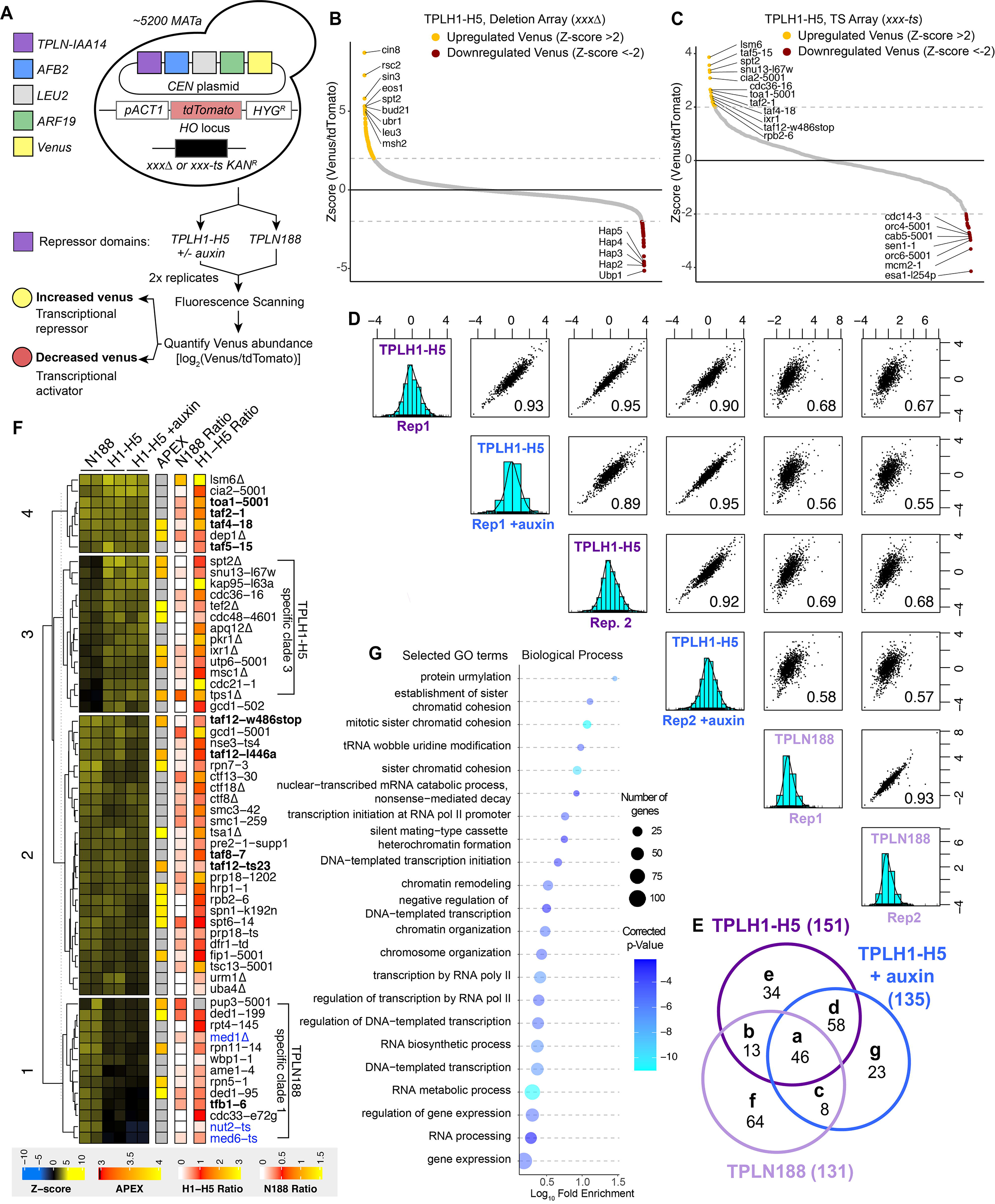
Reporter SGA identifies genetic interactors with TPL that are enriched for transcriptional machinery. A. Scheme for R-SGA screening to identify interactors that modulate TPL-based repression. This approach assayed a TPL repressed promoter driving Venus and RPL39pr-tdTomato abundance across ∼5200 yeast mutants carrying TPLH1-H5 repressor (+/- auxin) and TPLN188 repressor by fluorescence. **B.** Distribution of relative Venus abundances across all Deletion Array mutants screened with the TPLH1-H5 repressor. **C.** Distribution of relative Venus abundances across all Temperature Sensitive Array mutants screened with the TPLH1- H5 repressor. **D.** Correlation between all Temperature Sensitive Array experiments. Values at bottom right indicate the correlation value between biological replicates and conditions. The light blue graph indicates the distribution of z-scores in each sample. **E.** Venn diagram demonstrating the overlap between TPLH1-H5, TPLH1-H5 + auxin, and TPLN188 repressed screens for mutants with increased Venus abundance. These represent the intersection of results that were found in both biological replicates. **F.** Heat map of all TS screens and corresponding data from APEX and validation experiments. Columns 1-6 are z-scores for all mutants with upregulated Venus expression. APEX – log enrichment value from proximity labelling, gray – not detected. N188 ratio - independent cytometry validation of upregulated Venus mutant strains grown in liquid culture. H1-H5 ratio - independent cytometry validation of upregulated Venus mutant strains grown in liquid culture. Mutants in general transcription factors are highlighted in bold, mutants in mediator complex components are highlighted in blue. Tree was k- means clustered to highlight specific clusters of mutants (numbers on left). **G.** Selected significant biological processes present among mutants with increased Venus abundance organized by Gene Ontology (GO) terms.

To conduct the R-SGA, a strain containing the SPARC and expressing tdTomato constitutively from the *ACT1* promoter was mated with both the non-essential gene deletion array (DA) and conditional temperature-sensitive (TS) essential gene mutant collections using the SGA methodology^44^. We performed this approach with two different versions of the SPARC that differed only in the lengths of TPL N-terminal domains. The first SPARC contained helices 1-5 (SPARC^H1-H5^), where the only repressor domain is the LisH domain (H1-H2), and the second contained the entire N-terminal domain (SPARC^N1^^88^) comprised of helices 1-9 (H1-H9) and carries both the LisH and a second repressor domain in Helix 8, which contacts the Med21 subunit of the Mediator complex^21^. Each experiment was performed with two biological replicates, resulting in 4 collections each containing ∼ 5200 strains harboring a SPARC, pACT1- tdTomato and either a unique nonessential gene deletion (*xxx*Δ) or a TS essential gene allele (*xxx-TS*). Using a fluorescence scanner, we assayed the Venus and tdTomato intensities across the mutant strains arrayed as whole colonies on agar plates, and then we calculated a normalized log2(Venus/tdTomato) ratio, which is indicative of Venus abundance. For collections with SPARC-TPL^H1-H5^ the screen was also tested on media containing auxin, as this truncation is within the dynamic range for auxin sensitivity (Figure 2A).

We hypothesized that mutants causing increased Venus expression in the absence of exogenous perturbation (untreated conditions) would unveil two classes of genes: (1) those upregulating Venus transcription and therefore candidates for partner proteins in transcriptional repression and (2) those that downregulate Venus expression and likely involved in the activation ARF transcription (Figure 2A). By further performing the SPARC R-SGA screen on two isoforms of TPL, we aimed to identify genetic factors specific to the LisH domain of unknown molecular mechanism. Testing the R-SGA in the presence of auxin should reduce the cellular concentration of the repressor, increase Venus signal output for all mutants, and help to differentiate specific hits from background effects. Our focus in this study was on the first class of genes, those where genetic interaction yielded a “loss of repression” phenotype, as these represent the best TPL pathway candidates.

We applied a Z-score based threshold to identify mutants with the greatest change in Venus abundance, defining those mutants with Z > 2 (corresponding to a Venus abundance two standard deviations above the mean) as having increased Venus and those with Z < −2 as having decreased Venus. Using these thresholds, in collections with SPARC^H1-H5^ (TS plus DA) we identified 162 mutants (3% of those screened) with increased Venus abundance in untreated conditions and 56 with decreased Venus abundance (1% of those screened, Figure 2B-C,E, and Supplemental Tables 1,2). In SPARC H1-H5 on auxin treatment, we identified 147 mutants (2.8% of those screened) with increased Venus abundance and 67 with decreased Venus abundance (1.3% of those screened, Supplemental Tables 1,2). The screen with SPARC^TPLN1^^88^ identified 141 mutants (2.65% of those screened) with increased Venus abundance and 45 with decreased Venus abundance (0.85% of those screened, Supplemental Tables 1,2). A larger fraction of mutants with increased Venus abundance were nonessential (∼67%); however, a greater relative proportion of all essential mutants screened had increased Venus abundance (e.g., ∼32% of the 230 mutants with increased Venus are essential, although essential genes comprise only ∼19% of the mutants screened). The trend for higher genetic interaction in essential genes is consistent with the high number of core transcriptional machinery in the TS collection.

The average correlation across replicate screens was R = 0.94 and R = 0.83 for the temperature-sensitive collection and nonessential deletion collection, respectively. To graphically highlight the TS as an example, the average correlation within the SPARC^H1-H5^ was R = 0.92, and within SPARC^TPLN1^^88^ was R = 0.93, indicating strong reproducibility across biological replicates (Figure 2D, DA replication Figure S2). These correlations are similar to previous R-SGA screens, where an average of R = 0.77 was observed in technical replicates across 27 screens^43^. We further honed our target list by eliminating frequently found mutations called “frequent fliers” that have been discovered in previous studies to be non-specific hits in many R-SGA analysis^45^. We then focused our attention on mutants where we observed an increase in Venus abundance, and compared these hits between repressor types and conditions (Figure 2E). We observed 46 unique mutants that are shared between all repressor types (Figure 2E, group a), as well as mutants that were enriched only in specific repressors or conditions (Figure 2E, groups e, f and g).

To best highlight categories of mutants, we performed hierarchical clustering analysis of upregulated mutants that met our criteria (TS - Figure 2F, DA – Figure S3). Within the TS array, we observed three distinct clusters: (1) mutants identified in both repressor types (Figure 2F, clusters 2 & 4), (2) mutants identified specifically in the TPLN188 repressor (cluster 1) and (3) mutants identified only the H1-H5 repressor background (cluster 3). One clear and exciting observation is that the TPLN188-specific cluster 1 was the only cluster containing Mediator mutants (Figure 2F, blue) and is consistent with the observation that TPL binds to Mediator through its H8 repressor domain, which is absent in the H1-H5 truncation. We also observed that across these clusters there were many mutants in GTF genes (Figure 2F, bold), and at a broad level using GO analysis we could see that the mutants identified from both arrays are enriched for genes involved in transcription, especially initiation and regulation of RNA Pol II gene expression (Figure 2G). To validate upregulated mutants for a specific effect on SPARC repression, we selected the top 242 upregulated mutants for secondary analysis by cytometry. In this way, we were able to validate a large proportion of mutants in liquid culture (H1-H5: DA - 74%, TS - 80%, N188: DA - 57%, TS - 69%, Figure S4, Figure S5), and confirm that many top candidates are increased in Venus fluorescence with limited changes in cell size (Figure S6). Because we observed that ScSpt5 was highly enriched in TPL-APEX2 labeling and because we detected other Spt-phenotype related genes through R-SGA (i.e. ScSpt2, ScSpt6, ScSpt21, ScHTA1/Spt11, ScSpn1), we repeated R-SGA strain construction for the TS mutant *spt5-194* as it had failed to sporulate well in R-SGA conditions. Similar to the other SPT strains, we observed a strong de-repression of both the H1-H5 and N188 SPARCs in the *spt5-194* mutant (Figure S4-6).

### TPL interacts with the SPT4/SPT5/SPT6 complex through the conserved elongation factor SPT4

To find the overlap between physical and genetic interactors, we compared our list of 531 APEX2-labeled nuclear localized proteins against genes identified by R-SGA (Figure 3A, teal, Figure S7). Based upon previous studies we expected a low overlap between these two methods^46^. For example, only 0.5% of positive genetic interaction pairs from a genome-wide network also showed a protein–protein interaction^46^. We observed that 52 genes (9%) were found by both assays: 40 up-regulated (Figure 3A, yellow) and 12 down-regulated (Figure 3A, red). We focused on the 41 genes that are both up-regulated in R-SGA and detected by proximity labeling. We next ranked these candidates by how strongly enriched these proteins were by APEX labeling. To do so, we compiled our list of 40 overlapping interactors onto the 531 nuclear labeled proteins by their APEX2 enrichment relative to our known protein interaction partners (ARF, AFB, TPL). This approach identified five proteins that were enriched at a level equal to or greater than that of TPL itself: ScSpt5, ScTef4, ScRpb2, ScCdc48, and ScDed1.

**Figure 3.**
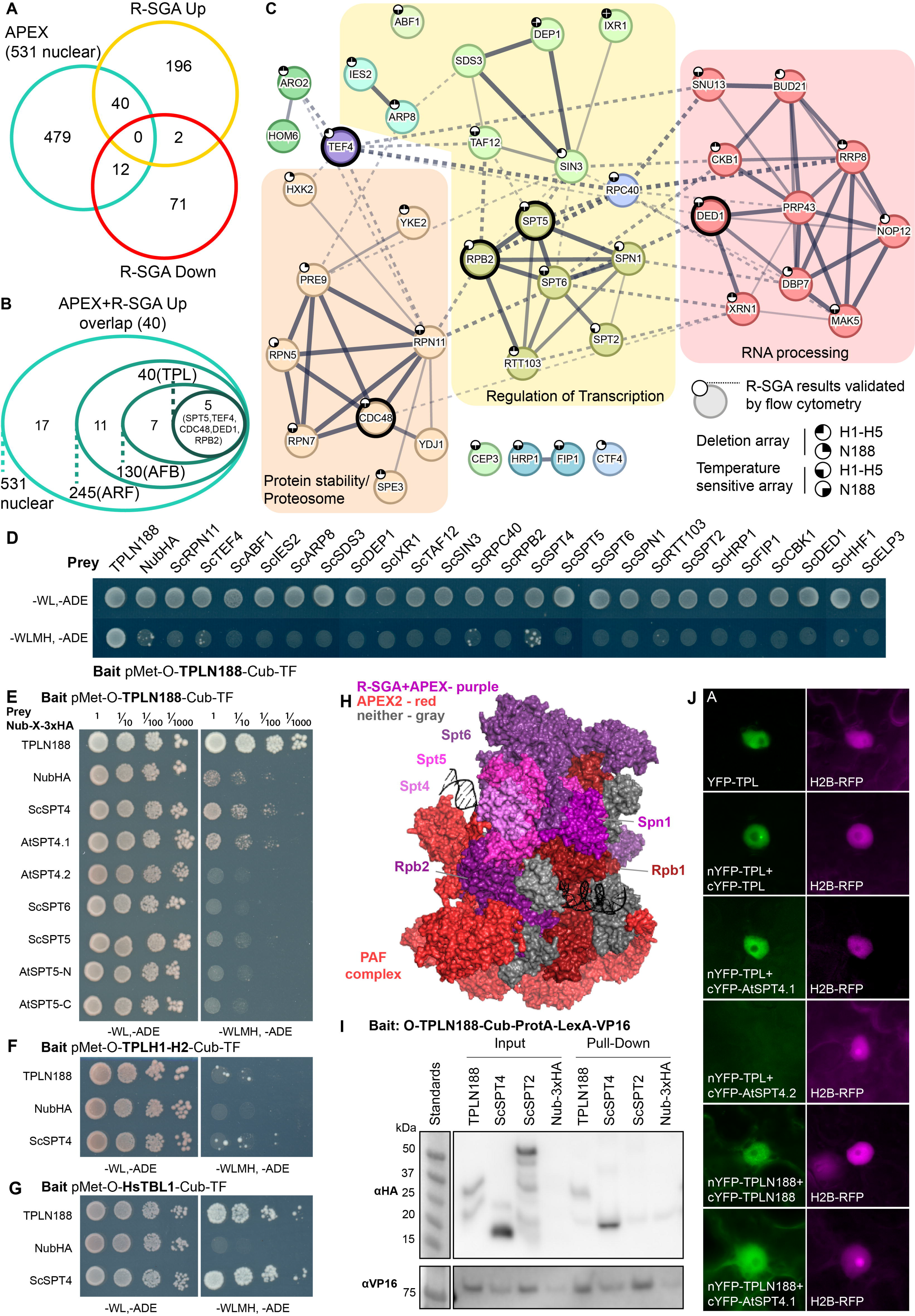
TPL interacts with the SPT4/SPT5/SPT6 complex through the conserved elongation factor SPT4 **A.** Venn diagram intersection of genes identified by APEX labelling and R-SGA analysis. 531 nuclear proteins from APEX (teal), 238 up-regulated R-SGA mutants (gold), and 85 down-regulated (red) mutants. **B.** A nested circle diagram depicts the APEX enrichment of the 40 proteins from the overlapping results of APEX and Up regulated R-SGA. Nested circles represent the relative enrichment positions of known controls (TF-ARF, F-Box-AFB2, TPLN188). Five proteins were enriched at or above the level of the TPLN188-TPLN188 interaction (SPT5, TEF4, RPB2, CDC48, and DED1). **C.** STRING protein network of the top 40 interacting proteins. The network is enriched for three main categories as defined by GO terminology: proteasome/protein stability - orange, regulation of transcription - yellow, and RNA processing - red. The five most highly enriched by APEX-labelling are highlighted with a heavy weighed outline. The network was clustered to a MCL inflation parameter of 3. Subclusters of genes are colored, and network edges between clusters are shown as dotted lines. Line thickness indicates the strength of support. STRING minimum required interaction score – 0.4. Small circles at the top left of the node indicate whether this R-SGA mutant was validated by flow cytometry, and in which TPL-truncation it was validated. **D-G.** Cytoplasmic split-ubiquitin system (CytoSUS) assays with candidate interacting proteins. Nub-3xHA is the N-terminal fragment of ubiquitin expressed with no fusion protein and is used as a negative control. -WL, -ADE: dropout lacking Trp, Leu, and Ade (growth control); -WLMH, -ADE: dropout lacking Trp, Leu, His, Met, and Ade (selective media). The plating for each panel was performed at the same day. **D.** CytoSUS interaction of TPLN188 with yeast proteins identified in the top 40, all individuals enriched for GO terms relating to transcription were tested in addition to other selected candidates. **E.** CytoSUS interaction of TPLN188 with SPT/DSIF specific components from yeast and *Arabidopsis*. **F**. CytoSUS interaction of TPL H1-H2 (LisH domain) with ScSPT4. **G.** CytoSUS interaction of HsTBL1 N-terminal domain (N98) with ScSPT4. **H.** Structure of RNA POL II subunits, DNA, PAF complex, with SPT/DSIF proteins – from yeast (*komagataella phaffii*) 7XN7^30^. Proteins identified by both APEX2 labelling and R- SGA - purples (SPT5, SPT6, RPB2, SPN1), identified only by APEX2 – reds, undetected – gray, SPT4 – pink. **I**. The TPL-ProteinA-TF fusion protein can pull down TPL, ScSPT4 from yeast extracts using IgG-beads. Detection of the VP16 transcriptional activator demonstrates enrichment of the fusion protein (αVP16). Each prey protein is detected via the 3xHA tag (αHA). **J**. Bimolecular fluorescence complementation assay performed in *Nicotiana benthamiana* (tobacco). Each image is an epi-fluorescent micrograph taken at identical magnification from epidermal cells two days post injection. The YFP image is colored green (left panel). p35S:H2B-RFP was used as a control and is false-colored magenta (right panel).

We generated a protein network of the 40 overlapping proteins that illustrates the known protein interactions between each of the proteins from the STRING database. A majority of genes in this network were also independently validated by flow cytometry analysis of the R-SGA identified mutants (Figure 3C, small circles). We observed three broad categories based on GO analysis (Figure 3C): proteasome/protein stability (orange), regulation of transcription (yellow), and RNA processing or stability (red). We observed a significant interconnected network within the group of genes with functions related to transcription. Specifically, we observed a highly interconnected cluster of genes comprised of ScRpb2, ScSpt5, ScSpt6, ScSpn1, ScSpt2, and ScRtt103 (Figure 3C, bolded outline).

To test whether these top hits are direct interactors with the TPL N-terminus, we introduced these *Saccharomyces cerevisiae* genes into the yeast cytoplasmic split-ubiquitin system (cytoSUS)^47^. We tested 24 of the 40 top hits comprising all of the proteins related to transcriptional regulation and several other representative selections from other clusters. We observed that none of these proteins had a strong direct physical interaction by cytoSUS, however, SPT4 was observed to interact with TPL (Figure 3D, Figure S8A). SPT4 and SPT5 are strong binding partners that co-purify across eukaryotic species^26,48^, and both *Saccharomyces* (ScSpt4) and *Arabidopsis* (AtSPT4.1) homologs interacted with TPLN188 (Figure 3E, Figure S8B). Similar to the yeast homologs, we observed no direct interaction between TPLN188 and AtSPT5 or AtSPT6. AtSPT4.2 failed to interact with TPLN188. Sequence divergence between the paralogs is highest in the C-terminus, likely pointing to the region of interaction with TPL (Figure S8C).

TPLN188 interacts with Med21 through the CRA domain, specifically H8^21^, whereas the H1 repressor domain has no currently defined interactors. We observed that ScSpt4 interacted with bait constructs containing only the TPL LisH domain (H1- H2) (Figure 3F). The human protein Transducin Beta-like 1 (HsTBL1) contains a LisH domain with a similar structural fold as TPL^49^, and we have previously demonstrated that the HsTBL1 N-terminal domain can repress transcription in yeast^22^. Like TPLN188, the HsTBL1 N-terminal domain strongly interacted with ScSpt4 (Figure 3G). These results indicate that TPL makes contact with Spt4 through the LisH domain.

A visualization of the PIC protein complex including RNA POL II subunits, DNA, PAF complex, and SPT or DSIF proteins^30^ highlights the density of proteins identified by APEX alone (red) or by both APEX2 and R-SGA (purple; ScSpt5, ScSpt6, ScRpb2, ScSpn1) (Figure 3H). SPT4 (pink) was not identified by proximity labeling or genetic screen. SPT4’s small size may have limited labeling or detection, and no *spt4* mutant is included in the SGA mutant collection. To further validate the interaction between SPT4 and TPL, we performed co-immunoprecipitation between TPLN188 and ScSPT4 in yeast (Co-IP; Figure 3I), and bimolecular fluorescence complementation (BiFC) between full-length TPL and AtSPT4.1 in plants (Figure 3J). In both assays, we observed a clear interaction between TPL and SPT4.

### The SPT4/SPT5/SPT6 complex is functionally required by TPL for repression

TPLN188 interacted with SPT4 through its LisH domain (Figure 3, 4A), and we hypothesized that it was through this interaction that it controls the activity of the SPT complex containing SPT4, SPT5 and SPT6. SPT5 and SPT6 loss of function mutants are lethal in yeast^50,51^, and the point mutations *spt5-194* (S324F^48^) and *spt6-14* (S952F^33^) we identified in the R-SGA screen library are temperature sensitive alleles. To test the requirement for SPT4, SPT5 and SPT6 in maintaining repression of a TPL- repressed promoter, we used the Anchor Away^52^ system for inducible protein depletion from the nucleus (Figure 4B) and combined it with quantification of transcriptional activity at the synthetic locus in the SPARC (Figure 2A). Through nuclear depletion of the SPTs, we could quantify transcriptional activity through increased fluorescence measured by flow cytometry^21^ (Figure 4B). We first tested the previously published ScSpt5^53^ and ScSpt6^54^ anchor away yeast strains for modulation of TPL repression of the SPARC. We observed release of repression through increased fluorescence for both ScSpt5 and ScSpt6 anchor away strains with both SPARC^N1^^88^ (Figure 4C) and SPARC^H1-H5^ (Figure 4D), but not for ScSpt2 anchor away (Figure S9A). Importantly, we observed that the ScSpt5 or ScSpt6 anchor away more potently released repression in TPL H1-H5, and there was no contribution to repression from ScMed21, which we have previously demonstrated is specific to H8 (Figure 4D).

**Figure 4.**
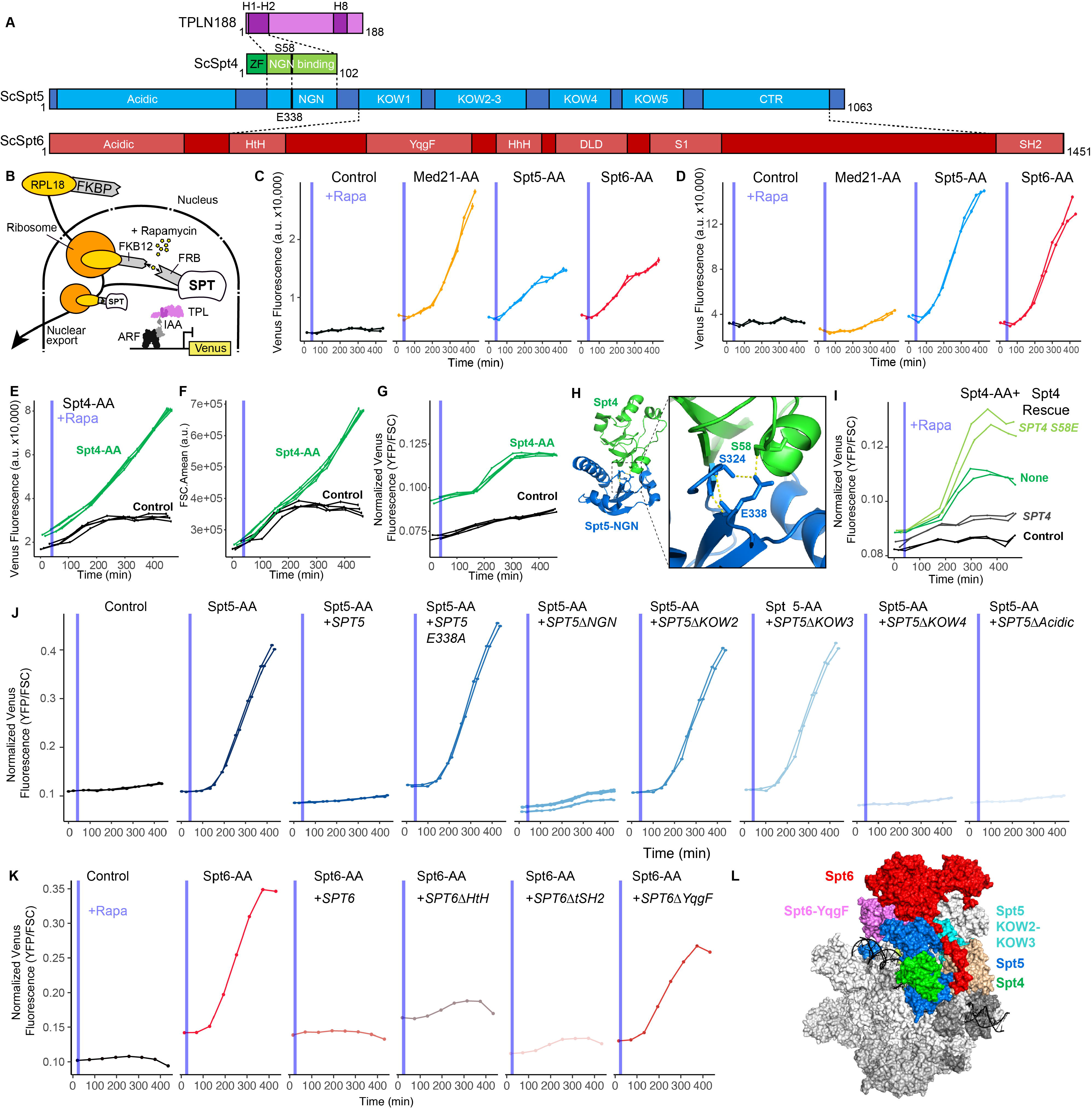
The SPT4/SPT5/SPT6 is functionally required by TPL for repression **A.** A schematic of the annotated domains of *Arabidopsis thaliana* TPLN188, and *Saccharomyces cerevisiae* Spt4, Spt5, and Spt6. Dotted lines indicate known protein interactions between these proteins. The position of the known interaction residues of SPT4 and SPT5 are shown with a black line and residue numbering. **B.** Schematic of inducible expression and nuclear depletion of SPTs in the context of the Auxin response circuit (ARC). In Anchor away, the yeast ribosomal protein 13A (RPL13A) is fused to the rapamycin-binding protein FKBP. Additional of rapamycin induces dimerization between FKBP and any target protein fused to 2xFRB, resulting in removal of the target protein from the nucleus. Expression of the ARC is monitored by expression of Venus fluorescent protein. C-G,I-K. Each panel represents representative data from at least two independent time-course flow cytometry experiments of the specific conditions indicated. Every point represents the average fluorescence of 5–10,000 individually measured yeast cells (a.u.: arbitrary units). Rapamycin (Rapa -1 µM) was added at the indicated time (gray bar, +Aux). **C.** Time-course flow cytometry analysis of SPARC^N188^ in Med21, Spt5 and Spt6 Anchor Away strains. **D.** Time-course flow cytometry analysis of SPARC^H1-H5^ in Med21, Spt5 and Spt6 Anchor Away strains. **E-G.** Time-course flow cytometry analysis of SPARC^H1-H5^ in Spt4 Anchor Away strains. E – Raw fluorescence (FL2.Amean), F – Cell size (FSC.Amean), G – Normalized fluorescence (FL2.Amean/ FSC.Amean). **H.** Structure of the SPT4-SPT5 binding interface PDB:2EXU^48^. Inset – zoom in on the acid dipole interaction interface with critical residues labelled and hydrogen bonds shown in yellow. **I.** Time-course flow cytometry analysis of SPARC^H1-H5^ in Spt4 Anchor Away strains with selected genome-integrated SPT4 rescue constructs. **J.** Time-course flow cytometry analysis of SPARC^H1-H5^ in Spt5 Anchor Away strains with selected genome-integrated *SPT5* rescue constructs. **K.** Time-course flow cytometry analysis of SPARC^H1-H5^ in Spt6 Anchor Away strains with selected genome-integrated *SPT6* rescue constructs. **L.** Structure of RNA POL II subunits, DNA, PAF complex, with SPT/DSIF proteins – 7XN7^30^. SPT6 – red, SPT6-YqgF domain – pink, SPT5 – blue, SPT5-KOW2-KOW3 – teal, SPT4 – green.

SPT5 and SPT6 have well defined activity in the context of their physical interactions with RNA Pol II^55–57^, and SPT4 interacts with SPT5 through its NGN-binding domain^48^ (Figure 4A). We created SPT4 anchor away strains to test whether SPT4 was required for TPL-based repression. Indeed, fluorescence levels increased upon ScSpt4 anchor away (Figure 4E); however, we also observed increases in cell size (Figure 4F), which has been well characterized in ScSpt4 loss of function mutations in yeast where *spt4-*Δ is among the largest 5% of haploid deletion strains^58^. After normalizing for cell size, we observed that loss of ScSpt4 does lead to a modest release of TPL repression (Figure 4G). The SPT4-SPT5 binding interaction is coordinated by an acid-dipole interaction directed by Ser58 in SPT4 and Glu338 in SPT5 (Figure 4A,H)^48^. To test whether ScSpt4 is required for repression without relying on a full loss of function, we performed a genetic rescue experiment by introducing either wild type ScSpt4 or a Ser58 mutant (S58E) into our ScSpt4 anchor away strain. We observed complete rescue with wild type ScSpt4 in the presence of Rapamycin. ScSpt4^S58E^ was completely unable to rescue repression despite retaining nearly normal cell size (Figure S9B-D), suggesting that the SPT4-SPT5 contact is required for repression (Figure 4I).

On the SPT5 side of the SPT4-SPT5 interaction face, the *spt5-194* mutation (S324F) identified in R-SGA likely destabilizes hydrogen bonding between Ser324 and Glu338, introduces a bulky residue into the binding pocket, and is therefore likely to destabilize SPT4-SPT5 interaction^48^ (Figure 4H). To interrogate whether reciprocal mutations in ScSpt5 that disrupt ScSpt4 binding affect repression, we created a deletion of the NGN domain and a Glu338 to alanine mutation in ScSpt5 rescue plasmids. We observed that the NGN domain deletion had a dramatic effect on cell size, and also largely eliminated reporter signal, similar to the ScSpt4 anchor away results (Figure 4J, Figure S9E). However, the E338A mutation was more specific, phenocopying the ScSpt5 anchor away, providing further support for a model where SPT4 and SPT5 together are required for H1-mediated TPL-based repression (Figure 4J).

Our observation that loss of SPT4, SPT5 and SPT6 activity resulted in a loss of repression led us to hypothesize that some complexes containing these components must be required for this repressive function. Both SPT5 and SPT6 are multidomain proteins that make extensive contacts with both RNA Pol II and other complexes^26,27^. We engineered targeted deletions in both SPT5 and SPT6 in selected domains to determine which may be required for this newly found repressive function (Figure 4A,J,K). We observed that in ScSpt5 KOW2 and KOW3 domains are required for repression, but not the acidic or KOW4 domains (Figure 4J, Figure S9E). In ScSpt6, the YqgF domain was required for repression, but not the tSH2 domain (Figure 4K, Figure S9F). The YqgF domain has recently been implicated in initiation of transcription, as mutant forms lacking this domain are trapped at the TSS in *Arabidopsis*^37^, while the tSH2 domain is well documented as being a Pol-II interaction surface^27^. We then mapped the location of these domains onto the protein complex formed between RNA POL II subunits, DNA, PAF complex, and SPT or DSIF proteins as in Figure 4H^30^.

Strikingly, the KOW2 and KOW3 domains in ScSpt5 and the YqgF in ScSpt6 are each found at the junction between these proteins and RNA Pol II subunits RPB1 and RPB2 (Figure 4L).

### TPL requires components of the nucleation step of the transcriptional pre-initiation complex (PIC) for repression

If Mediator and SPT4-6 transcriptional regulators are required for repression, it is logical to wonder whether other components of the general transcription machinery are similarly involved. In the first stages of promoter identification, the TATA box binding protein (TBP), a subunit of the TFIID complex, binds to the promoter at the TATA box and induces DNA bending (Figure 5A). TBP then recruits TFIIA and then TFIIB to the promoter, which in turn recruits RNA Pol II and TFIIF. Finally, TFIIE is recruited and brings TFIIH, with kinase activity, to phosphorylate the RNA Pol-II CTD, at which point the PIC is transcriptionally active. The TFIID complex was highly enriched by APEX proximity labeling, as were subunits of TFIIA and TFIIE (Figure 5A, red and purple). We also observed multiple subunits from TFIID in our R-SGA data, as genetic interactors with the SPARC (Figure 5A, blue and purple). In the context of the TFIID structure^59^, the extensive physical and genetic interaction with TAF subunits became even more apparent (Figure 5B).

**Figure 5.**
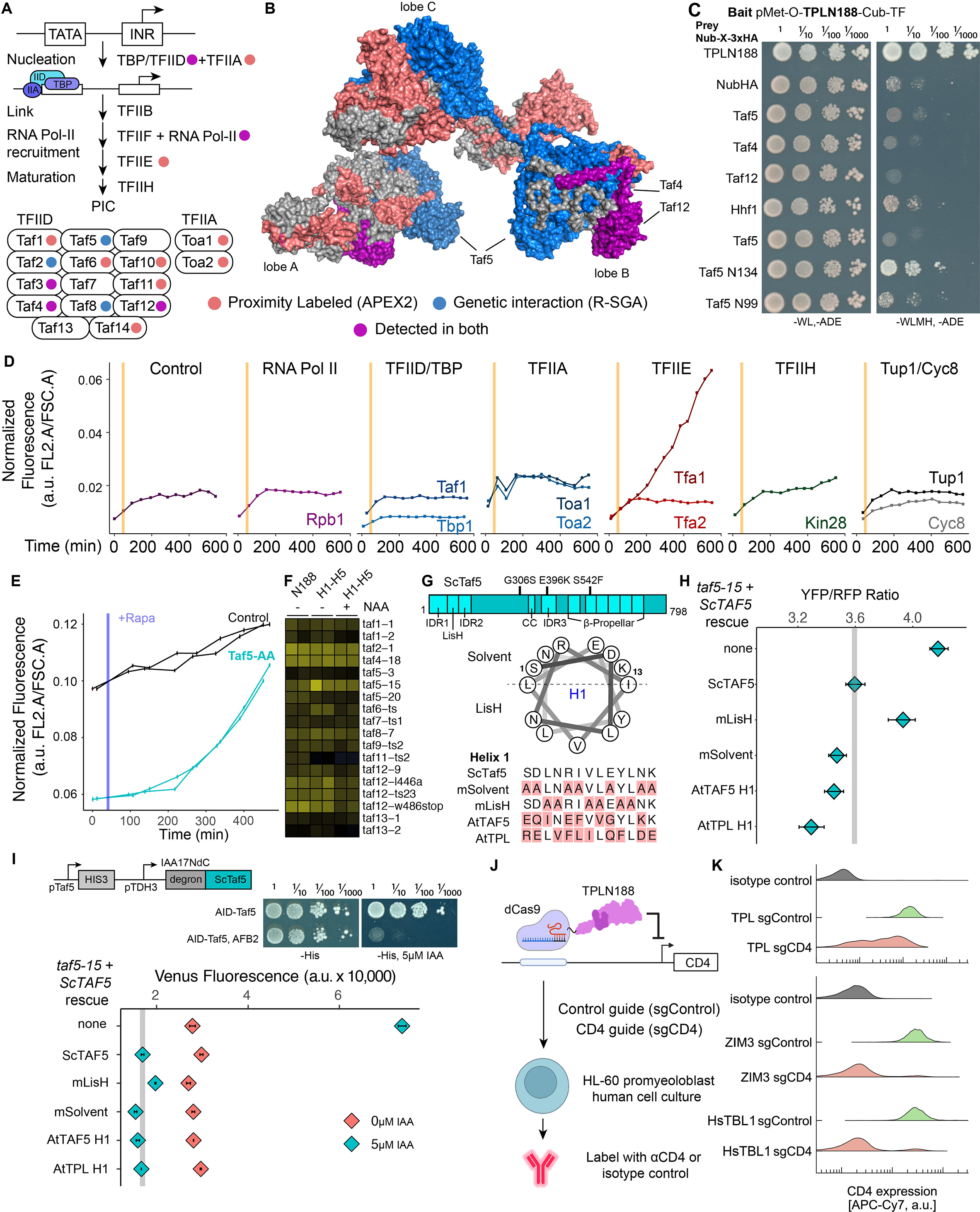
TPL requires components of the nucleation step of the transcriptional pre-initiation complex (PIC) for repression. A. Schematic of the steps of the transcriptional pre-initiation complex formation on a promoter, accompanied by a cartoon representation of TFIID and TFIIA components **B.** Crystal structure of human TFIID PDB:6MZL^59^. Both A and B were colored to represent which genes were identified by proximity labelling (red) and genetic interaction (blue) or both (purple). **C.** Cytoplasmic split-ubiquitin system (CytoSUS) assays with candidate interacting proteins. Nub-3xHA is the N-terminal fragment of ubiquitin expressed with no fusion protein and is used as a negative control. -WL, -ADE: dropout lacking Trp, Leu, and Ade (growth control); -WLMH, -ADE: dropout lacking Trp, Leu, His, Met, and Ade (selective media). **D.** Time-course flow cytometry analysis of SPARC^H1-H5^ in selected Anchor Away strains that include selected GTF components^63^. **E.** Time-course flow cytometry analysis of SPARC^H1-H5^ in a Taf5 Anchor Away strain (Taf5-2xFRB). **F.** Heat map of all TFIID components in TS screens. Columns 1-6 are z-scores for all mutants that were recovered from all TS replicate screens. NAA – auxin application. **G.** Top – gene map of the ScTaf5 protein. Middle – helical wheel diagram of the ScTaf5 LisH Helix 1 alpha helix. Bottom – protein alignment of ScTaf5 LisH Helix 1 domain and selected mutations. **H.** Steady state flow cytometry of *taf5-15* with selected rescue constructs grown at the non-permissive temperature (30°C). Gray bar indicates the fluorescence of the wild-type rescue construct. **I.** AID-Taf5 schematics and rescue experiments. Fluorescence were quantified by cytometry at 6 hours after application of 5μM IAA. **J.** Cartoon schematic of the dCAS9-repressor experiment to test LisH repression function in HL-60 cell culture. **K.** Quantification of CD4 protein levels by flow cytometry. Isotype control is provided to highlight the baseline fluorescence levels.

Previously, we identified that ScTaf5, the structural core of TFIID for both lobe A and B (Figure 5B) contains a LisH domain that was capable of repressing transcription in our synthetic yeast assay^22^. This led to our hypothesis that perhaps the TPL LisH domain could dimerize with the Taf5 LisH domain to recruit TPL to the TFIID complex at both A and B lobes (Figure 5B). However, we detected no interaction between TPL and full-length ScTaf5, ScTaf4, or ScTaf12 proteins. We did observe low level interactions between TPL and the Histone H4 protein (ScHhf1) as has been previously described^60^. This interaction is thought to be driven by the histone tail, so it is perhaps not surprising we did not see interaction with Taf4 or Taf12 through their histone fold domains (Figure 5C). However in the case of HsTAF5, it has been well characterized that the HsTAF5 WD40 domain requires the TRiC/CCT chaperone to fold and subsequently is rapidly recruited into the HsTAF5-6-9 complex^61,62^, and therefore may be physically unavailable to bind in the CytoSUS assay. Truncations of ScTaf5 containing only the N-terminal domain IDRs and LisH (N134) showed strong TPL binding activity, which is drastically reduced but not eliminated upon deletion of the LisH domain (N99, Figure 5C, Figure S10A).

To begin to test whether components of the PIC are required for repression, we used a previously characterized series of anchor away strains that include selected GTF components^63^. Unsurprisingly, depletion of proteins required for transcription such as ScRpb1 or ScTbp results in yeast that cannot transcribe any genes including the reporter (Figure 5E, Figure S10B). However, anchoring away TFIIE did abrogate transcriptional repression (Figure 5E, Tfa1). Both ScTbp and ScTaf1 anchor away strains led to a loss of growth (Figure S10B). Therefore, we engineered a ScTaf5 anchor away strain, which showed the elevation of reporter expression that is consistent with a role for TAF5 in repression (Figure 5E). However, it is clear that the C-terminal fusion of the 2xFRB tag required for anchor away reduced the function of ScTaf5, as reporter transcription prior to Rapamycin addition was quite low (Figure 5E).

To independently assess a role for TAF5 in TPL-mediated repression, we turned to the TFIID TS mutants identified by the R-SGA (Figure 5F). *taf5-15* (G306S,E396K,S542F, Figure 5G) had the strongest signal in R-SGA experiments and was selected for further experiments. To test which residues within the ScTaf5 LisH domain might be critical for TPL-mediated repression, we mapped the residues in ScTaf5 with respect to their predicted orientation in the context of the LisH dimerization interface for Helix 1 (Figure 5G, helical wheel). We engineered alanine mutations in the solvent facing (mSolvent), LisH interface (mLisH), as well as domain swaps where the LisH Helix 1 was replaced with the Helix 1 from either *Arabidopsis* TAF5 (AtTAF5) or TPL (AtTPL, Figure 5G). We introduced these constructs into *taf5-15* mutant strains carrying the SPARC^H1-H5^. Wild type ScTAF5 fully rescued repression, as did mSolvent, AtTAF5, and AtTPL (Figure 5H). In contrast, mLisH failed to rescue repression, indicating that the conserved residues in the H1 domain that direct LisH dimerization are required for the activity of ScTAF5 in its role in repression. This finding is consistent with cross-kingdom analyses of other LisH domains^22^. We engineered an Auxin inducible degron (AID) version of ScTaf5 (AID-Taf5, Figure 5I) where the degron was fused to the N-terminus of ScTaf5 and observed that this approach did not reduce function of the Taf5 protein (Figure 5I). We performed the same rescue experiments as in Figure 5H, and observed similar results, with the exception that AID-induced loss of ScTaf5 had an even greater effect on fluorescence levels (Figure 5I), and that mLisH exhibited a more modest effect on reporter output.

The TPL N-terminal domain has been structurally compared^49^ to other LisH domain containing proteins such as the Transducin-Beta like 1 (HsTBL1^64^), which is a component of the SMRT/NCoR complex^6^, and acts as an exchange factor, facilitating conversion of SMRT/NCoR repressed loci to transcriptionally active states^65^. TPL H1 and H1-H2 (the full LisH) are functional repression domains in yeast^22^, and our findings here pointed to a functional interaction repertoire that should be conserved across all eukaryotes. To further test this hypothesis, we introduced dCAS9-TPLN188 into human HL-60 cells and compared its repression capacity with the KRAB-domain containing protein domain from ZIM3, which recruits KAP1/TRIM28 to induce DNA-methylation^66^ (Figure 5J). In this experiment, the CD4 gene was targeted and protein levels were quantified by immunostaining and flow cytometry^67^. We observed a significant reduction in CD4 abundance when TPLN188 was recruited to the CD4 promoter region, compared to a control guide (Figure 5K). Interestingly, the HsTBL1 N-terminal domain (N98), which we previously found to be able to replace TPL in our synthetic yeast assay^22^, was equally efficient at repressing CD4 as ZIM3 (Figure 5K).

### Transcription by TPL *in planta* requires the LisH domain and its interacting partners SPT6 and TAF5

To test the impact of TPL interactors on repression in an endemic developmental context, we used a quantitative repression assay based on UAS/GAL4-VP16^21,68,69^ (Figure 6A-B). We generated transgenic *Arabidopsis* lines where the UAS-TPL- IAA14^mED^ constructs were activated in the xylem pole pericycle stem cells where IAA14 normally acts to regulate the initiation of lateral root primordia (Figure 6A). To block potentially confounding interactions with endogenous TPL/TPRs or TIR1/AFBs, we engineered a variant of IAA14 with mutation in the two EAR domains (EAR^AAA^) and in the degron (P306S, hereafter termed IAA14^mED^; Figure 6B). We created mutations in the TPL LisH-H1 sequence to investigate its repressive function in lateral root development (Figure 6C). Expression of functional TPLN188-IAA14 fusion proteins in these xylem pole pericycle cells strongly suppressed the production of lateral roots, phenocopying the solitary root (*slr*) mutant, which carries the auxin-resistant form of IAA14 (Figure 6D). Disrupting the wild-type TPL H1 sharply decreased repression and restored lateral root density (Figure 6D). We observed that the LisH was required for repression even when the H8 repression motif was intact (Figure 6D). This result suggests an order of operations to the TPL mechanism, where TPL interacts first with SPT4/TAF5 and then with MED21 to establish a durable repression state. Interestingly, mutations in either solvent or buried faces of the LisH rendered the TPL protein non-functional in *Arabidopsis* (Figure 6D), indicating that these both play a critical role in repression, marking a point of divergence between yeast and plants. Additionally, we uncovered a difference between ScTaf5 and AtTAF5, where the AtTAF5 has a degenerate LisH motif that is incapable of replacing the TPL H1 sequence (Figure 6C,D).

**Figure 6.**
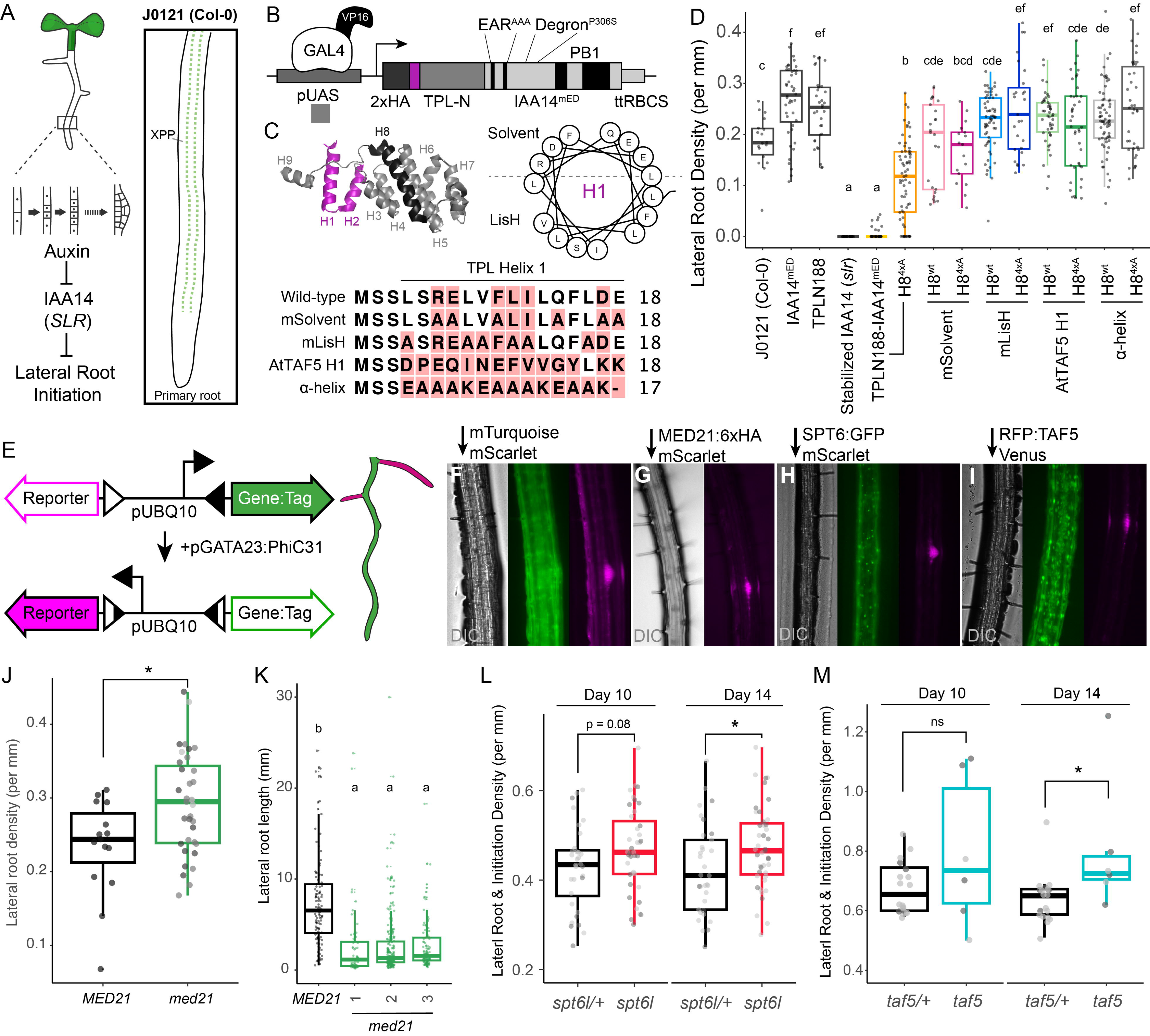
Transcription by TPL *in planta* requires the LisH domain and its interacting partners MED21, SPT6L and TAF5. **A.** Auxin-induced degradation of the IAA14 is strictly required for initiation of lateral root development (cartoon). An enhancer trap line (J0121) expresses GAL4-VP16 and UAS-GFP in the xylem pole pericycle cells. **B.** Design of UAS-TPL-IAA14^mED^ construct. Mutation of the conserved lysine residues in the EAR domain disrupted potential interactions with endogenous TPL/TPR proteins. The IAA14 degron has been mutated (P306S) to render it auxin insensitive. UAS: upstream activating sequence; ttRBCS: Rubisco terminator sequence. **C.** The protein structure of the N-terminal domain of AtTPLN188, with LisH domain highlighted in blue (left) Helical wheel depiction of AtTPL H1 sequence colored by their physicochemical class (Red/blue, charged; yellow, hydrophobic) (right) The consensus sequence of LisH-H1 sequence with different residue mutations in gray are displayed below. **D.** LisH mutations in TPL were sufficient to break repression of the development of lateral roots in Arabidopsis seedlings. The density of emerged lateral roots was measured in T1 seedlings at 14 days after germination. **E.** Design of the integrase target. The target is composed of two integrase sites (triangles) surrounding a constitutive promoter (pUBQ10) and two fluorescent reporters (GFP and mScarlet). In the absence of the integrase, GFP is expressed. In the presence of the integrase, there is an inversion of the DNA between the target sites, inverting the promoter to drive mScarlet expression. The expression of the integrase is mediated by the GATA23 promoter, which is expressed in the founder cells of lateral roots. **F-I**. Epifluorescence micrographs of roots of T1 plants from representative integrase switch lines. All images were taken at 12 days after germination at 20x magnification. **J**. The effect of loss of MED21 on lateral root density (emerged roots only) was calculated at day 14 in T2 plants. Data points from each independent line appear with the same grey value. **K**. Lateral root lengths at day 14 for plants in panel J. Letters indicate significant difference (ANOVA and Tukey HSD multiple comparison test; p<0.001). **L,M.** Lateral root initiations (emerged roots + non-emerged primordia) were quantified in heterozygous and homozygous mutant lines at day 10 and 14. Data points from each independent line appear with the same grey value. Asterisks indicate significant difference (ANOVA and Tukey HSD multiple comparison test; p<0.05).

We next wanted to test whether the SPTs and GTFs we identified in yeast play analogous roles with TPL corepressor activity in plants. However, like in yeast, many of these genes are essential in *Arabidopsis* (*AtSPT4*^70^, *AtSPT5*^70^, *AtSPT6L*^37^ & *AtTAF5*^71^). To develop a conditional loss of function assay, somewhat analogous to anchor away in yeast, we took advantage of recent work deploying serine integrases to mediate site-specific and irreversible DNA recombination in *Arabidopsis* roots^72^. This framework allowed us to engineer loss of gene function in a specific cell type during development (Figure 6E) by rescuing null T-DNA mutants with a constitutively expressed wild-type gene with a promoter flanked by integrase sites. When expression of the integrase PhiC31 is under the control of pGATA23, it triggers an irreversible inversion of DNA between integrase sites only in xylem pole pericycle cells. This integrase switch leads to a loss of expression of the essential gene and concomitant expression of a fluorescent reporter in those cells and all descendants.

We used this approach to study three target genes *AtMED21*, *AtSPT6L* and *AtTAF5* that met the criteria of being single copy genes and having available loss of function T-DNAs carrying glufosinate resistance. The latter condition made it possible to use characterized target and Integrase constructs. We selected homozygous mutant lines for each gene of interest, carrying both a target construct and pGATA23:PhiC31.

For each experiment, we identified lines with cell-type specific switching (Figures F-I) and quantified the effect of loss of candidate gene function on lateral root density (Figure 6J,L,M). For each TPL-interacting target gene, we observed an increase in root initiation events when the gene function was lost (Figure 6J,L,M), consistent with the prediction that loss of repression would mimic the accumulation of auxin and subsequent induction of IAA14-repressed loci. As expected from their essential role in transcription, the lateral roots produced in these lines were not fully wild-type. In all cases, the length of the lateral roots was sharply reduced. In the case of MED21, stubby roots emerged from the primary root, making it possible to measure their length compared with wild-type plants (Figure 6K). In roots where SPT6L and TAF5 function was lost, lateral root development was aborted before emergence, making it necessary to include earlier stages in our analysis (Figures L,M).

## Discussion

Over the last decade, advances in cryoEM^73^ and crystallography^74–76^ have provided an increasingly clear picture of the transcription Preinitiation Complex; however, corepressors are largely missing from these studies. Despite overlapping structural features and functions^12,23^, there is little agreement on how this class of regulatory proteins inhibit PIC activity^19,21,24,77^. Here, we investigated this question using a single synthetic locus paired with whole-genome screens for protein-protein (APEX2, Figure 1) and genetic interactors (R-SGA, Figure 2) with the corepressor TPL. We focused our efforts on the N-terminal domain of TPL for several reasons: (1) it is sufficient for repression in both yeast and in plants^20–22^, (2) its structure has been solved^42,49^, (3) it contains two discrete repression domains^21^, and (4) it has well-characterized contact points with other proteins in our synthetic circuit that can serve as internal controls (Aux/IAAs, MED21, MED10, and TPL itself^11,21,42,49^). The results here corroborate our previous work connecting one of TPL’s repression domains to the Mediator complex^21^, and build on these findings by linking the second repression domain to core PIC subunits. Specifically, we found that the LisH domain, found in the first two helices of TPL, recruits the DRB Sensitivity Inducing Factor (DSIF) complex components SPT4 and SPT5, the interacting protein SPT6, and likely Pol-II itself (Figure 3). Moreover, the interaction between TPL and the SPT genes was required to maintain repression in both yeast (Figure 4) and plants (Figure 6). In addition, general transcription factors in the TFIID complex were identified through both APEX and R- SGA, with TAF5 directly interacting with TPL to maintain repression (Figure 5). These results taken together suggest that TPL nucleates the PIC at repressed loci, while inhibiting progression of transcription until an activation signal has been received.

By acting as a bridge between sequence-specific and general transcription factors, corepressors and coactivators act at many loci to regulate whether a PIC is assembled and whether it is licensed to initiate transcription. Transcription initiation is a multi-step process starting with assembly of the TBP/TFIID and TFIIA complexes^74^. In the case of the synthetic TPL-regulated promoter used in these studies, we observed stable binding of the activating transcription factor (ARF19), the Aux/IAA adaptor protein and TPL, accompanied by SPT4, SPT5 and SPT6 (Figure 3, 4), elements of TFIIA & TFIID (Figure 2, 3, 5), and Mediator^21^. This complex was not sufficient to initiate transcription, although we did observe association of Pol-II subunits by proximity labeling. As we were unable to immunoprecipitate Pol-II at the synthetic promoter^21^, we hypothesize that the interaction of Pol-II with the repressed locus may be relatively weak or transient.

While SPT4, SPT5 and SPT6 are best known for their roles in transcriptional elongation^27,30,78–80^ and proximal promoter pausing in metazoans^81–83^, they were originally identified as yeast mutants that were able to restore transcription at a locus with a promoter disrupted by the Ty transposon^32^. The Suppressor of Ty genetic screen identified many core components of the transcriptional machinery, including: histones, TBP, Mediator15, as well as components of SAGA and FACT complexes^84^. The inclusion of SPT4, SPT5 and SPT6 in this group suggests a role in promoter identification and transcription initiation phases in addition to aiding passage of Pol II during elongation. Subsequent studies found that SPT5 stabilizes Pol II protein and regulates chromatin state at enhancers in human cell culture^85^. The bacterial homolog of SPT5, NusG, is negatively regulated by direct binding of the Rho protein, which triggers dynamic structural conformation changes in RNA Polymerase subunits and termination^86^. In *Salmonella*, depletion of NusG leads to massive upregulation of silenced sites that depend on this Rho-NusG mechanism of repression^87^. Recent work in *Arabidopsis* found that the YqgF domain of SPT6, the same domain we found was required for TPL-mediated repression, is crucial for the initiation of transcription^37^.

Repression of PIC formation at the level of TFIID is an effective mechanism to inhibit transcription of genes in euchromatin as has been demonstrated by work on several repressors (NC2 ^88,89^, MOT1^90^, CBF1^91^, adenovirus E1A^92^, Rb^93^). For example, the negative regulator NC2/Dr1-DRAP1 that acts on the TFIID/TBP step of initiation inhibits transcription initiation by Pol II via direct interactions with the TBP-DNA binary complex^94^. Within TFIID, the TAF5 protein has several characteristics that make it of particular interest. ScTaf5 shares a structural homology to corepressors, with a N-terminal LisH domain followed by WD40 beta-propeller domains (Figure 5G). AtTAF5 has lost the functional LisH domain, while other lineages (i.e. metazoan and fungal) have conserved this domain, and this may partially explain why the repression by TPLN188 is stronger in yeast than in plants^20,21^. The isolated LisH domain of yeast ScTaf5 represses transcription when targeted to chromatin^22^, indicating that an inherent repressive function has to be overridden during activation. Inter-LisH domain binding has also been observed in assembly of the GID E3 ligase complex, a multi-subunit E3 complex that achieves higher order assembly through LisH interactions^95^.

As TPL is functional as a transcriptional repressor in plant, yeast and human cells, an attractive model is that a variety of corepressors with similar structural features prime loci for rapid activation by facilitating pre-assembly of the PIC, including looping of sequence-specific transcription factors bound to enhancer elements to stabilize binding of TFIID, TFIIA and DSIF. By stabilizing a specific partially assembled state of the PIC (i.e. TFIID, SPTs, Mediator), TPL and other LisH-domain containing proteins such as the human HsTBL1 protein, may create a primed state that allows rapid entry into a transcription competent state. Indeed, TPL and HsTBL1 were sufficient to repress transcription in a synthetic context in human cell culture, highlighting the conservation of this mechanism of PIC inhibition across eukaryotes. The next challenge will be to map the dynamics and mechanism by which these primed complexes are disassembled allowing transcription to progress, and how the primed state is re-assembled once an activating signal is removed.

## Acknowledgments

We thank current and former members of the Nemhauser group including Cassandra Maranas, and Dr. Sarah Guiziou and for constructive discussions and comments on this manuscript; Dr. Adam Steinbrenner and Dr. Takato Imaizumi for insightful suggestions; Prof. Grant Brown for helping to lay the groundwork for this project during the 2018 yeast course at Cold Spring Harbor Labs; and Julie A. Theriot and Mathew Footer for supporting the experiments that relied on mammalian cell tissue culture. In addition, we thank Prof. Maitreya Dunham and Dr. Joe Armstrong for advice on yeast genetics and approaches; Prof. Brenda Andrews for donation of key R-SGA yeast strains; Prof. Joeseph C. Reese for donation of the SPT5 anchor away strains; Prof. Brian D. Strahl for donation of the SPT6 anchor away strains; Prof. Kevin Struhl for donation of the GTF anchor away strains; and Prof. Chen and Prof. Cui for donation of *Arabidopsis* SPT6L genetic constructs. We thank Daria Chrobok of DC SciArt for creation of the graphics adapted in this work.

This work was supported by the NIH (R01-GM107084 and R35-GM148135-01 to JLN; 5K99GM147355 to NMB; R35 GM119536 to JV), a Faculty Scholar Award from the Howard Hughes Medical Institute (to JLN), and the Canadian Institutes for Health Research (FDN-159913 to GWB). GWB holds a Canada Research Chair (Tier 1). Research support for NMB was also provided by the Howard Hughes Medical Institute. ARL was supported as a Simons Foundation Fellow of the Life Sciences Research Foundation.

## Author contributions

Conceptualization, Methodology, Project administration: ARL, JLN. Software – Programming: ARL. Validation – Verification, Formal analysis, Data Curation, Visualization: ARL, BD, JSS. Investigation: ARL, BD, JSS, RLK, NB, RARM, AB, IJW, LB. Resources - JV, GWB, JLN. Writing - Original Draft: ARL, BD, JSS, JLN. Writing - Review & Editing: ARL, BD, JSS, RLK, NB, RARM, AB, IJW, LB, JV, GWB, JLN. Supervision: ARL, JLN, GWB. Funding acquisition – JLN, GWB, JV.

## METHOD DETAILS

### Cloning

Construction of TPL-APEX fusion proteins was performed by Golden Gate cloning as described in Pierre-Jerome et al., 2014. Variant and deletion constructs were created using PCR-mediated site-directed mutagenesis. Construction of the SPARC was described previously^21^. Site-directed mutagenesis primers were designed using NEBasechanger and implemented through Q5 Site-Directed Mutagenesis (NEB, Cat #E0554S). For the cytoplasmic split-ubiquitin protein-protein interaction system, bait and prey constructs were created using the plasmids pMetOYC and pNX32, respectively (Addgene, https://www.addgene.org/Christopher_Grefen/). Interaction between bait and prey proteins was evaluated using a modified version of the split ubiquitin technique (Asseck and Grefen, 2018). TPL interactor genes were amplified either from yeast gDNA or as cDNAs from wild-type Col-0 RNA using reverse transcriptase (SuperScript IV Reverse Transcriptase, Invitrogen) and gene-specific primers from IDT (Coralville, IA), followed by amplification with Q5 polymerase (NEB). These cDNAs were subsequently cloned into plasmids for cytoSUS using a Gibson approach^96^. The coding sequence of the genes of interest was confirmed by sequencing (Genewiz; South Plainfield, NJ). For UAS-driven constructs, the TPLN188-IAA14 coding sequence was amplified with primers containing engineered BsaI sites and introduced into the pGII backbone with the UAS promoter and RBSC terminator^97^ using Golden Gate cloning as described previously^21^. Subsequent mutations were performed on this backbone using PCR-mediated site-directed mutagenesis (see above). Construction of C-terminal 2xFRB fusions for Anchor Away was done as described^52^. For construction of the Integrase target vectors the sequence of the rescue gene was amplified by PCR from template DNA with BsaI adaptors, and assembled into an integrase target vector as previously described^72^.

### Proximity labeling and protein preparation

Proximity labeling was performed according to previously published protocols for proximity-dependent proteomic profiling in yeast cells by APEX2 and Alk-Ph probe^38,39^. The general protocol was identical to these protocols with the following adaptations starting from step 27 of the protocol after reduction and alkylation (step 26): Beads with bound protein samples were washed twice with 1mL 100mM ammonium bicarbonate, resuspended in 200µL 100mM ammonium bicarbonate and digested with sequencing grade trypsin by incubating at 37°C for 16 hours. Digestion was stopped by addition of 10% TFA to a final concentration of 1% TFA. Precipitate was removed by centrifugation, peptides were loaded onto conditioned MCX extraction cartridges (Waters, Milford, MA). Peptides were washed with i) 0.1% TFA and ii) 75% ACN 0.25% FA. Clean peptides were eluted with 75% ACN, 2.5% NH4OH and dried down by vacuum centrifugation.

### Mass Spectrometry

Lyophilized peptide samples were resuspended in 3% ACN, 5% formic acid and subjected to liquid chromatography coupled to tandem mass spectrometry (LC-MS/MS). Samples were loaded into a 100 μm ID x 3 cm precolumn packed with Reprosil C18 1.9 μm, 120Å particles (Dr. Maisch). Peptides were eluted over a 100 μm ID x 30 cm analytical column packed with the same material housed in a column heater set to 50°C and separated by gradient elutionof 3 to 28% B (A: 0.15% FA, B: ACN 80% and 0.15% FA) over 60 min and 28% to 45% B over 13 min at 350 nl/min delivered by an Easy1200 nLC system (Thermo Fisher Scientific) with a total 90 min method length.

Peptides were online analyzed on a Orbitrap Eclipse Tribrid mass spectrometer (Thermo Fisher Scientific). Mass spectra were collected using a data dependent acquisition method. For each cycle a full MS scan (375-1500 m/z, resolution 120,000, and standard 100% normalized AGC target in automatic maximum injection time mode) was followed by MS/MS scans (isolation width 1.6 Da, 30% normalized collision energy, 30,000 resolution and standard 100% normalized AGC target in automatic maximum injection time mode) on the topmost intense precursor peaks with a 3s cycle time between full MS scans.

### Mass Spectrometry Data Analysis

Raw files were converted to the mzXML format using MSConvert (Proteowizard)^98^ and MS/MS spectra were searched against a target/decoy protein sequence database using Comet (version 2019.01.02)^99^. *Saccharomyces cerevisiae* (orf_trans_all.fasta downloaded from the Saccharomyces Genome Database in 2016) with the four protein components of the AtARC^Sc^ (AtARF19, AtAFB2, AtTPLN188-APEX2, AtIAA14) was used as database and search parameters tolerances chosen based on Comet recommendations for high resolution MS1 and MS2 acquisitions (i.e. 20 ppm precursor mass tolerance, 0.02 Da fragment tolerance for MS/MS acquired). Trypsin was selected as the digestive enzyme with a maximum of 2 missed cleavages, fixed carbamidomethylation modification of cysteines (+57.0215 Da) and variable modifications of methionine oxidation (+15.9949 Da) and protein N-terminal acetylation (+42.0106 Da). Search results were filtered with Percolator^100^ (version 3.01) to 1% false discovery rate at the peptide spectrum match level. Peptide abundance was determined using in-house quantification software to extract MS1 intensity. Protein Prophet^101^ was used to assemble peptides into protein groups and roll up peptide quantifications into protein quantifications.

### Yeast Methods

Standard yeast drop-out and yeast extract–peptone–dextrose plus adenine (YPAD) media were used, with care taken to use the same batch of synthetic complete (SC) media for related experiments. A standard lithium acetate protocol^102^ was used for transformations of DNA. All cultures were grown at 30°C with shaking at 220 rpm.

Anchor Away approaches were followed as described^52^, and Anchor Away strains were obtained from EUROSCARF (euroscarf.de). Endogenous genomic fusions of GENE- FRB were designed by fusing gene homology to the pFA6a-FRB-KanMX6 plasmid for chromosomal integration into the parental Anchor Away strain, selectable through G418 resistance (G418, Geneticin, Thermo Fisher Scientific). Tup1-FRB and Cyc8-FRB were constructed as described^21^. Mediator and GTF Anchor Away strains were created previoulsy^63^ and kindly donated by Dr. Kevin Struhl. SPT5^53^ and SPT6^54^ anchor away yeast strains were previously published. SPARC construction was described previously^21^. For the cytoplasmic split-ubiquitin protein-protein interaction system, bait and prey constructs were created using the plasmids pMetOYC and pNX32, respectively (Addgene, https://www.addgene.org/Christopher_Grefen/). Interaction between bait and prey proteins was evaluated using a modified version of the split ubiquitin technique^103^. After 2 days of growth on control and selection plates, images were taken using a flatbed scanner (Epson America, Long Beach, CA).

### SPARC R-SGA

Two independent crosses of YNL3669 (*MATα SPARC^H1-H5^[pTDH3-AtTPLH1-5-IAA14-ttACS, pRPS2-AtAFB2-ttCIT1, LEU2, pADH1-AtARF19-ttADH1, pP3(2x)-UbiVenus-ttCYC1*] *HO::ACT1pr-tdTomato::hphMX, can1*Δ*::STE2pr-Sphis5 lyp1*Δ *his3*Δ*1 leu2*Δ*0 ura3*Δ*0 met15*Δ*0*) and YNL3670 (*MATα SPARC^TPLN1^*^88^*[pTDH3-AtTPLN188-IAA14-ttACS, pRPS2-AtAFB2-ttCIT1, LEU2, pADH1-AtARF19-ttADH1, pP3(2x)-UbiVenus-ttCYC1] HO::ACT1pr-tdTomato::hphMX, can1*Δ*::STE2pr-Sphis5 lyp1*Δ *his3*Δ*1 leu2*Δ*0 ura3*Δ*0 met15*Δ*0*) with the yeast nonessential deletion collection and a set of conditional temperature-sensitive alleles of essential genes were performed following standard SGA procedures^104^. Final arrays were pinned in duplicate on either SD/MSG–his–leu+ 200mg/mL G418 (untreated) or YPD supplemented with 50mM NAA and grown for 24hr before fluorescence scanning. The Typhoon TrioVariable Mode Imager (GEHealthcare) was used to acquire Venus (488- nmlaser, 520/40BP emission filter) and tdTomato (532-nmlaser, 610/30BPemission filter) fluorescence values. For the essential temperature-sensitive mutants, all growth was conducted at 23°C until the final growth before imaging, where they were grown at 30°C. After fluorescence imaging, colony size data were acquired by individually photographing plates with a Canon PowerShotG 24.0 megapixel digital camera using Remote Capture software. Data analysis followed essentially what is described in Kainth et al. (2009)^43^, with small variations. To summarize, background-subtracted yellow fluorescent protein (Venus) and tdTomato intensities were computed for each colony from .GEL images using GenePixPro version 7.0 software. Colony size was imaged on SPimager from S&P Robotics, Inc, and size information was calculated from individual photographs SGAtools. Border colonies, small colonies (colony area < 500pixels), and colony size information was calculated from individual photographs.

Border colonies, small colonies (colony area < 500pixels), and colonies with aberrantly low tdTomato values (bottom 0.05%) were removed before further analysis. log2(Venus/tdTomato) values were calculated and LOESS normalized for each plate. Using the log2(Venus /tdTomato) ratio as a metric for Venus abundance has the advantage that dividing by tdTomato corrects for any colony size dependent intensity effects. Finally, normalized log2(Venus/tdTomato) values were averaged across all replicate experiments and a Z-score calculated (See Supporting Information). All analyses were performed in R.

### Flow Cytometry

Fluorescence measurements were taken using a Becton Dickinson (BD) special order cytometer with a 514 nm laser exciting fluorescence that is cut off at 525 nm prior to photomultiplier tube collection (BD, Franklin Lakes, NJ). For validation of R-SGA strains by flow cytometry fluorescence measurements were taken using a Sony SA3800 spectral cell analyzer with 488 nm, 405 nm & 561 nm lasers exciting fluorescence (Sony Biotechnology, San Jose, CA). Events were annotated, subset to singlet yeast using the FlowTime R package^105^. A total of 10,000–20,000 events above a 400,000 FSC-H threshold (to exclude debris) were collected for each sample and data exported as FCS 3.0 files for processing using the flowCore R software package and custom R scripts^106,107^ (available in Github: https://github.com/achillobator/TPL-H1_Mechanism). Data from at least two independent replicates were combined and plotted in R (ggplots2).

### Anchor Away

Three days before running Anchor Away assay, cultures were struck out onto new plates with the proper dropout selection and grown at 30°C for 2 days. Single colonies were diluted into media at ∼1 event/microliter, grown overnight to ∼100 events/microliter, and aliquoted into 96 well deep-well culture plates^106,107^. Rapamycin was added to a final concentration of 1μM and samples were analyzed over a time course and tested approximately once per hour.

### CytoSUS

Diploid yeast cultures were grown overnight in CSM media with proper dropout selection. Overnight diploid yeast cultures were analyzed for OD_600_ with a biophotometer. All cultures were diluted to 1 OD, following this series dilutions of 1/10, 1/100, and 1/1000 were made for all cultures. All serial dilutions of diploid yeast cultures were plates reciprocally on all CSM selection plates in organized rows utilizing reverse pipetting to ensure uniform cultures. Plates were left open to evaporate excess liquid from cultures and incubated at 30°C. After 2 days of growth on control and selection plates, images were taken using a flatbed scanner (Epson America, Long Beach, CA).

### Western Blot

Yeast cultures grown overnight in SC media were diluted to OD_600_ = 0.6 and incubated until cultures reached OD_600_ ∼1. Cells were harvested by centrifugation, lysed by vortexing for 5 min in the presence of 200 µl of 0.5 mm diameter acid washed glass beads and 200 µl SUMEB buffer (1% SDS, 8 M urea, 10 mM MOPS, pH 6.8, 10 mM EDTA, 0.01% bromophenol blue, 1 mM PMSF) per 1 OD unit of original culture. Lysates were then incubated at 65° for 10 min and cleared by centrifugation prior to electrophoresis and blotting. Antibodies: anti-HA-HRP (REF-12013819001, Clone 3F10, Roche/Millipore Sigma, St. Louis, MO), anti-FLAG (F3165, Monoclonal ANTI-FLAG M2, Millipore Sigma, St. Louis, MO), anti-FRB (ALX-215-065-1, Enzo Life Sciences, Farmingdale, NY; Haruki et al., 2008), anti-VP16 (1-21) (sc-7545, Santa Cruz Biotechnology, Dallas TX), anti-GFP (ab290, AbCam, Cambridge, UK), anti-MYC (71d10, 2278S, Cell Signaling, Danvers, MA), and anti-PGK1 (ab113687, AbCam).

### Co-Immunoprecipitation

Co-IP from yeast was performed using the cytoSUS strains. Cultures were grown to OD_600_ 0.5 (∼1E7 cells/ml) using selective media, harvested, and resuspended in 200 μl extraction buffer (50mM Tris-HCL pH 8, 100mM NaCl, 5mM MgCl_2_, 1mM EDTA, 1mMDTT, 0.5mM PMSF, 10% Gylcerol, 0.25% NP-40) with protease inhibitors. Cells were lysed by vortexing 3 × 1 min full speed with 100 μl of 0.5 mm Acid Washed Glass Beads, clarified by centrifugation (1 min, 1000 rpm), and supernatant was mixed with 1 ml IP buffer (15 mM Na_2_HPO_4_, mw 142; 150 mM NaCl, mw 58; 2% Triton X-100, 0.1% SDS, 0.5% DOC, 10 mM EDTA, 0.02% NaN_3_) with protease inhibitors and incubated with 100 μl of IgG sepharose at 25°C for 2 hr with rotation. The beads were washed 1× with IP buffer and 2× with IP-wash buffer (50 mM NaCl, mw58; 10 mM TRIS, mw 121; 0.02% NaN_3_) with protease inhibitors. Protein was eluted with 50 μl of SUME (1% SDS, 8 M urea, 10 mM MOPS, pH 6.8, 10 mM EDTA) buffer +0.005% bromophenol blue by incubation at 65°C for 10 min and run on handmade 12% acrylamide SDS-PAGE gels, and western blotted accordingly.

### Bimolecular Florescence Complementation

Agrobacterium-mediated transient transformation of *N. benthamiana* was performed as previously described^108^. BiFC experiments were performed on 3-week-old *N. benthamiana* plants grown at 22°C under long days (16 hr light/8 hr dark) on soil (Sunshine #4 mix) as per Martin et al., 2009. pSITE vectors^109^ were used to generate BiFC constructs for MED21, SPT4, TPL, and TPLH8QuadA – proteins. In all cases, the combinations are N-terminal fusions of either the nEYFP or cEYFP to the cDNA of MED21 or TPL. RFP fused Histone H2B was used as a nuclear marker^110^. Injection of Agrobacterium strains into tobacco leaves was performed as in Goodin et al., 2002, but the OD600 of the Agrobacterium culture used was adjusted to 0.5. Two days after transfection, plant leaves were imaged using an epifluorescence microscope (Leica Biosystems, model: DMI 3000B).

### CRISPRi cell line construction

The dCa9-ZIM3, dCa9-TPLN188, dCas9-HsTBL1, and sgRNAs constructs were integrated into HL-60 cells using a lentivirus spinoculation protocol^67^. Lentiviral vectors containing dCas9 and sgRNA were produced in HEK293 cells. Transfected HEK293 cells were lysed to collect lentivirus and concentrated. Briefly, lentivirus was added to 1LmL cells (1L×L10^6 cells/mL) and polybrene reagent (final concentration of 1 μg/mL) in 24-well tissue culture plates. Cells were spun at 1000Lg for 2 h at 33 °C. Virus was removed and cells were placed in an incubator for 2 days prior to antibiotic selection for 6 days (dCas9: blasticidin 10 μg/mL; sgRNA constructs: puromycin 1 μg/mL).

### Immunolabeling for flow cytometry

Live-cell immunofluorescence measurements of cell surface expression for CD4 was performed on a Sony SH800 Cell Sorter. All staining and washes were done with cells suspended in phosphate-buffered saline (PBS) containing 2% hiFBS (heat-inactivated fetal bovine serum) and 0.1% sodium azide, chilled on ice. For each sample, one million cells were first resuspended in100 μL buffer containing 5 μL Fc Receptor Blocking Solution (Biolegend, #422302) and incubated for 15Lminutes. Cells were then spun down and resuspended in 100 μL of buffer containing fluorescently conjugated antibodies for one hour (5 μL of each antibody per sample): APC-Cy7 Mouse Anti-Human CD4 (1:20 dilution; BD Biosciences, #557871), APC-Cy7 Mouse IgG1, κ Isotype Control (1:20 dilution; BD Biosciences, #557873). Following staining, the samples were washed three times by resuspending in 300 μL of fresh buffer, chilled on ice. Following collection of flow cytometry data,.fcs files were exported and processed using the Python package FlowCytometryTools^111^ (v. 0.5.1).

### Plant Growth

For *Arabidopsis thaliana* experiments using the GAL4-UAS system (Laplaze et al., 2005), J0121 was introgressed eight times into Col-0 accession from the C24 accession and rigorously checked to ensure root growth was comparable to Col-0 before use.

UAS-TPL-IAA14mED constructs were introduced to J0121 introgression lines by floral dip method^112^. T1 seedlings were selected on 0.5× LS (Caisson Laboratories, Smithfield, UT)+ 25 µg/ml Hygromycin B (company) + 0.8% phytoagar (Plantmedia; Dublin, OH). Plates were stratified for 2 days, exposed to light for 6 hr, and then grown in the dark for 3 days following a modification of the method of Harrison et al., 2006.

Hygromycin-resistant seedlings were identified by their long hypocotyl, enlarged green leaves, and long root. Transformants were transferred by hand to fresh 0.5× LS plates + 0.8% Bacto agar (Thermo Fisher Scientific) and grown vertically for 14 days at 22°C. Plates were scanned on a flatbed scanner (Epson America, Long Beach, CA) at day 14. *slr* seeds were obtained from the Arabidopsis Biological Resource Center (Columbus, OH). For integrase switch experiments T2 plant lines harboring T-DNAs for either MED21 (*med21-1*, WiscDsLox461-464K13), SPT6L (*spt6l-7*, SAIL_59_G06) and TAF5 (*taf5-4*, SAIL_274_A04) were transformed with the floral dip method to generate integrase target lines, and then used to introduce each integrase construct into these established target lines. For T1 selection: 120Lmg of T1 seeds (∼2000 seeds) were sterilized using 70% ethanol and 0.05% Triton-X-100 and then washed using 95% ethanol. Seeds were resuspended in 0.1% agarose and spread onto 0.5X LS Bacto selection plates, using 25 µg/mL of kanamycin for target lines and 25 µg/mL kanamycin and 25 µg/mL hygromycin for lines with both the integrase and the target. The plates were stratified at 4 °C for 48 h then light pulsed for 6 h and covered for 48 h. They were then grown for 4–5 days. To select transformants, tall seedlings with long roots and a vibrant green color were picked from the selection plate with sterilized tweezers and transferred to a new 0.5X LS Phyto agar plate for characterization.

### Quantification and statistical analysis

All quantification and statistical analyses were performed in R, and the corresponding code has been deposited into GitHub: https://github.com/achillobator/TPL-H1_Mechanism.

---

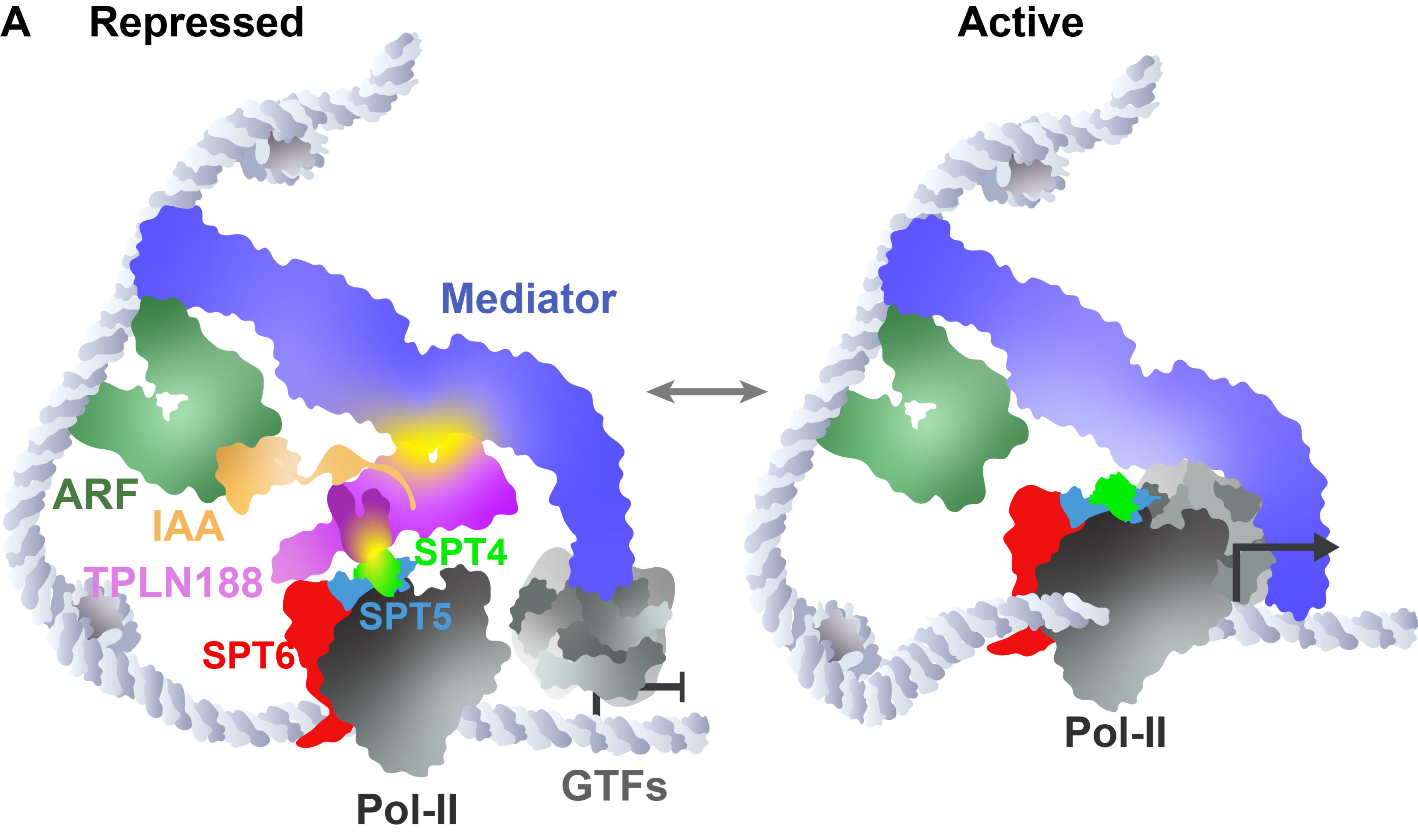

