## Supplemental Figures for "A conserved function of corepressors is to nucleate assembly of the transcriptional preinitiation complex"

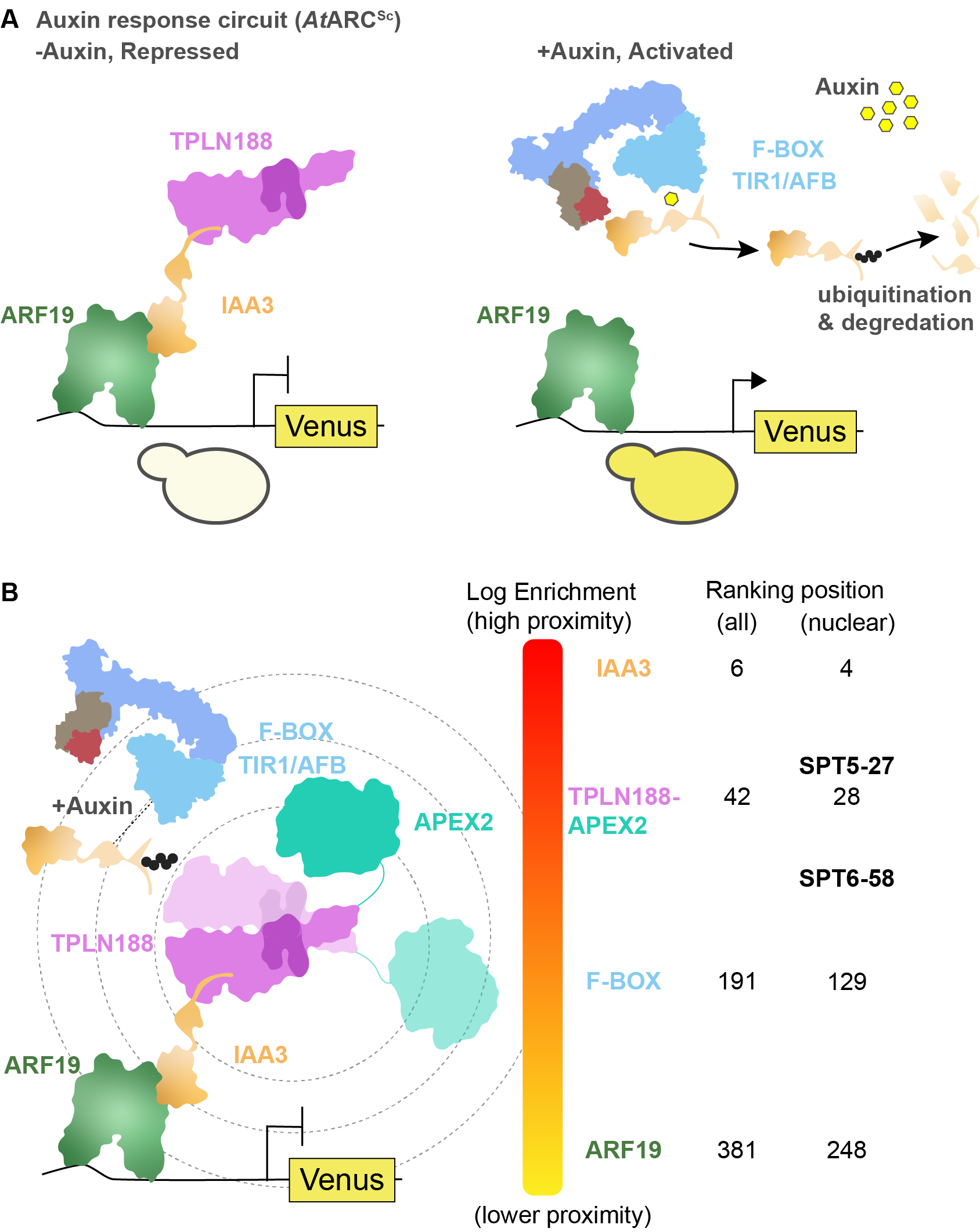

**Figure S1. A.** Schematic of the *At*ARC*^Sc^*. The auxin-responsive promoter driving the fluorescent protein Venus carries binding sites for the auxin-responsive transcription factor (ARF, here *AtARF19*). In the absence of auxin, the IAA protein (here *AtIAA3*) is bound to the TPL-N protein and to the ARF and maintains the circuit in a repressed state. The IAA protein can be used as a protein fusion with TPL, i.e. TPLN188-IAA3, or as separate parts as optimized in Figure 1. Upon addition of auxin, the IAA protein is targeted for ubiquitination and subsequent protein degradation, activating transcription of the fluorescent reporter. **B.** Proximity labelling benchmarked by relative enrichment compared to control protein interactors. Log2 fold enrichment values were benchmarked against the known experimental controls: AtTPLN188, AtIAA3, F-Box (AtAFB2 auxin receptor) and the ARF19 transcription factor. This ranking was calculated before and after subsetting for nuclear localized proteins (right column, determined from SGD). The relative ranking of ScSpt5 and ScSpt6 are shown in the nuclear column.

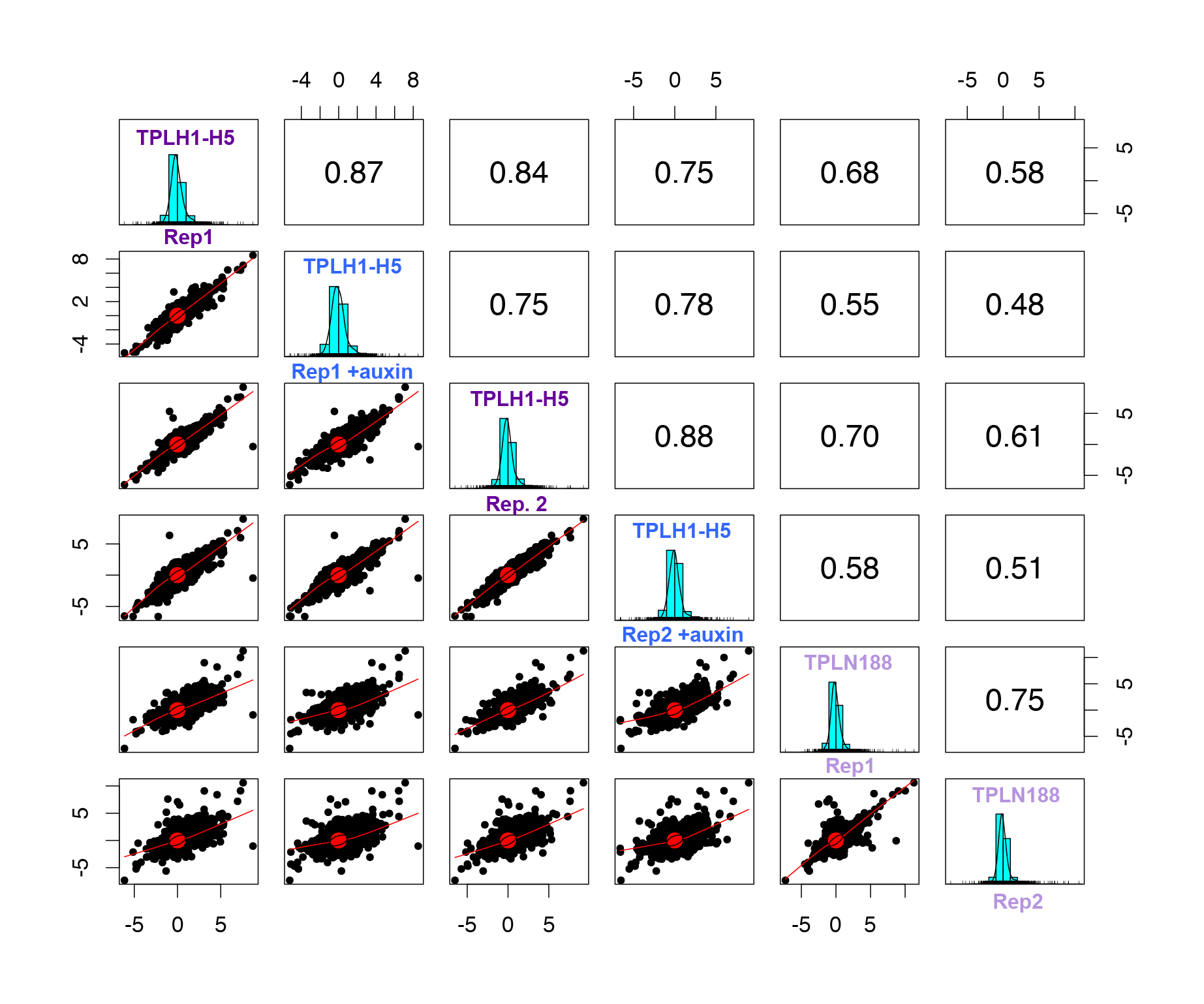

**Figure S2**. Correlation between all Deletion Array experiments. Values on the right indicate the correlation value between biological replicates and conditions. The light blue graph indicates the distribution of z-scores in each sample.

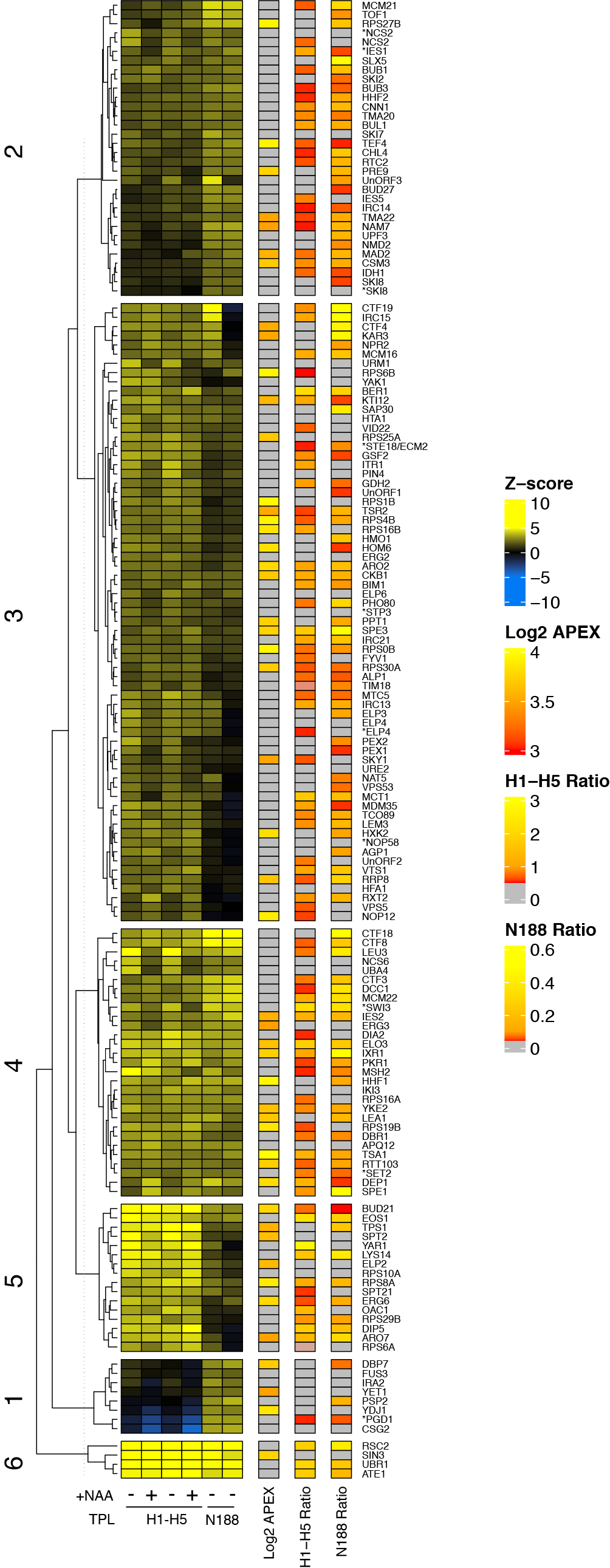

**Figure S3.** Distribution of relative Venus abundances across all Deletion Array mutants screened with the TPLH1-H5 repressor. Heat map of all DA screens and corresponding data from APEX and validation experiments. Columns 1-6 are z-scores for all DA mutants with upregulated Venus expression. APEX – log enrichment value from proximity labelling, gray – not detected. N188 ratio - independent cytometry validation of upregulated Venus mutant strains grown in liquid culture. H1-H5 ratio - independent cytometry validation of upregulated Venus mutant strains grown in liquid culture. Mutants in general transcription factors are highlighted in bold, mutants in mediator complex components are highlighted in blue. Tree was kmeans clustered to highlight specific clusters of mutants (numbers on left).

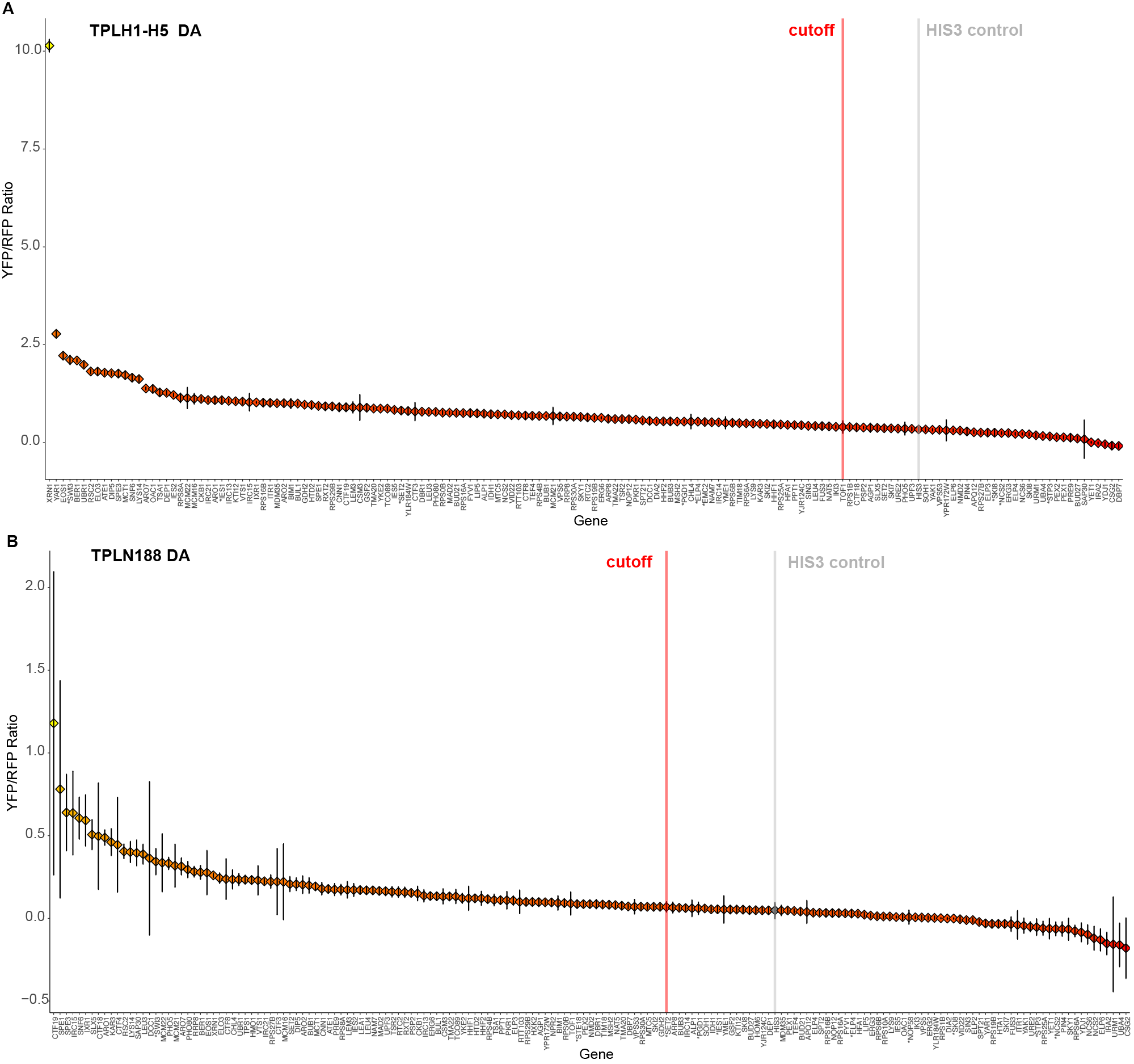

**Figure S4**. Cytometry validation DA. Selected strains were grown in liquid culture to log phase at 30°C and cytometry was performed. Fluorescence of tdTomato and Venus were quantified and Venus was normalized to tdTomato and is presented here as a ratio. Each data point is colored coded on a gradient with higher Venus expression as more yellow. Error bars are standard error propagated through the calculation of the ratio (error = sqrt((((((Venus.Asd/(sqrt(events)))/Venus.Amedian)^2) + (((tdtomato3.Asd/(sqrt(events)))/tdtomato3.Amedian)^2))*abs(Ratio)))). The cutoff was arbitrarily set as the value of control (grey line) plus its standard error (red line).

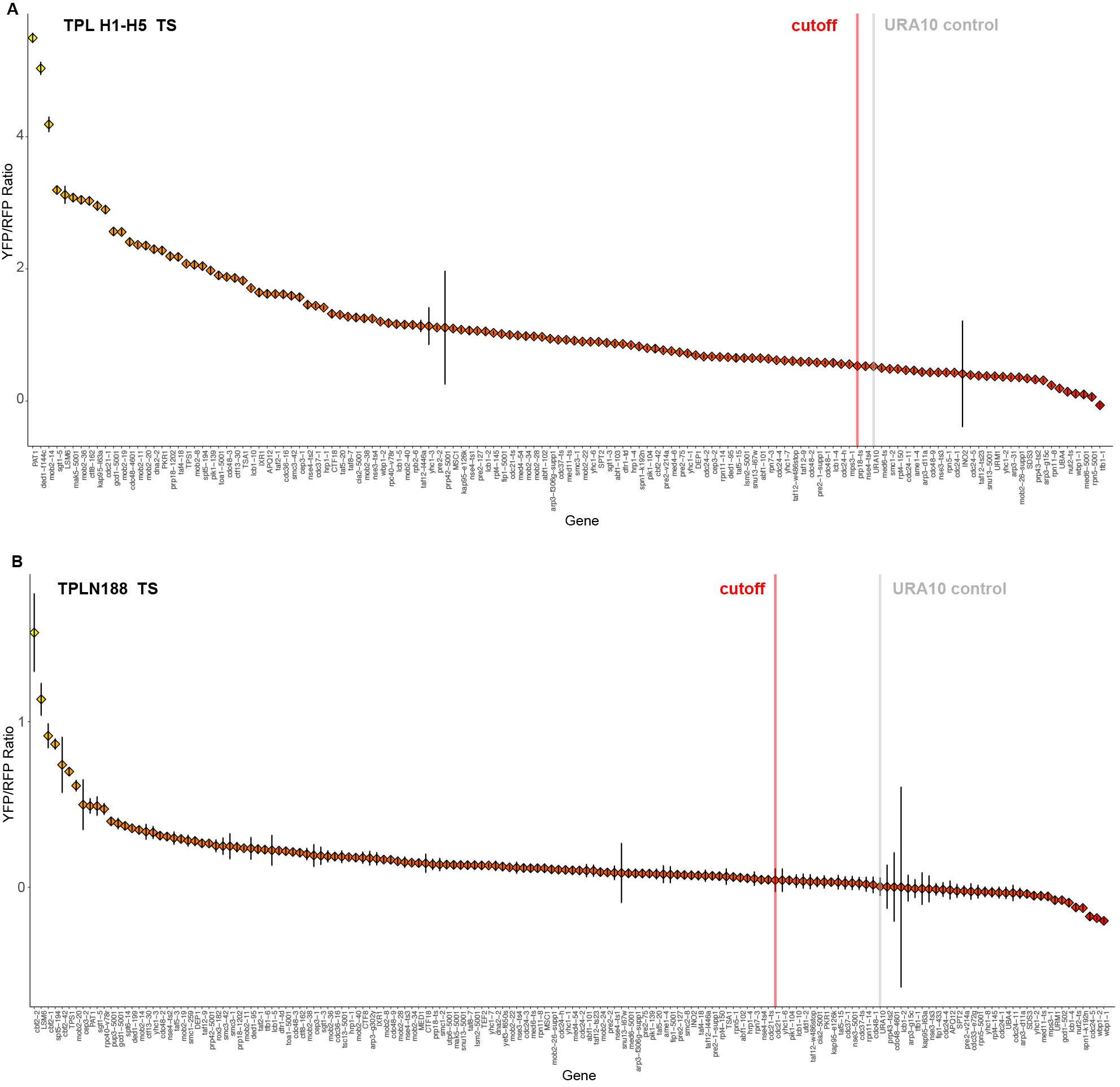

**Figure S5.** TS array Cytometry Validation. Selected strains were grown in liquid culture to log phase at 30°C and cytometry was performed. Fluorescence of tdTomato and Venus were quantified and Venus was normalized to tdTomato and is presented here as a ratio Each data point is colored coded on a gradient with higher Venus expression as more yellow. Error bars are standard error propagated through the calculation of the ratio (error = sqrt((((((Venus.Asd/(sqrt(events)))/Venus.Amedian)^2) + (((tdtomato3.Asd/(sqrt(events)))/tdtomato3.Amedian)^2))*abs(Ratio)))). The cutoff was arbitrarily set as the value of control (grey line) plus its standard error (red line).

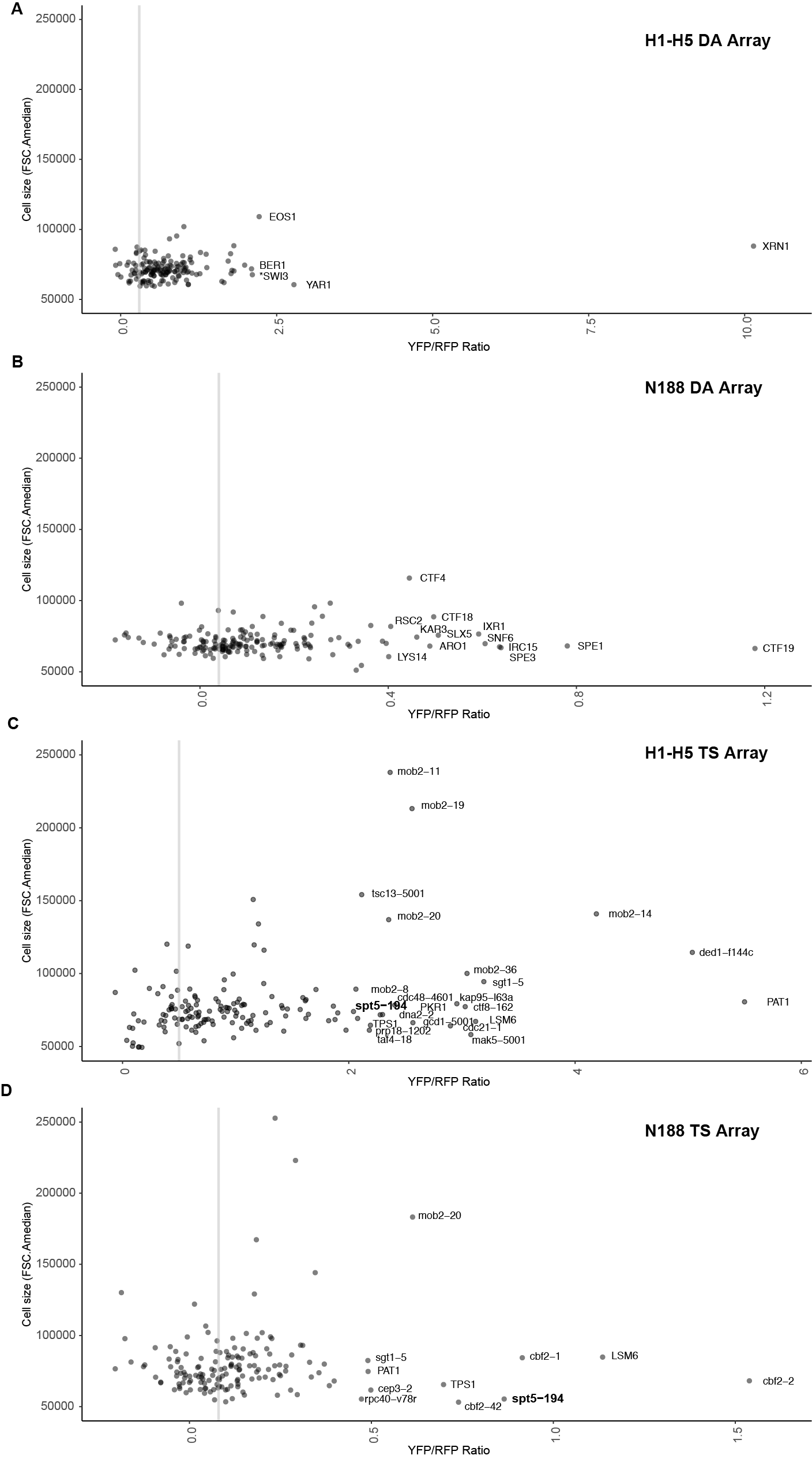

**Figure S6**. Yeast cell sizes from top hits cytometry validation. All cytometry validation experiments were plotted based on cell size (FSC.A) compared to their YFP/RFP ratios to highlight which candidates are likely to exhibit higher fluorescence values based on morphological differences. A. TPL H1-H5 deletion array. B. TPL H1-H5 TS array. A. TPL N188 deletion array. B. TPL N188 TS array.

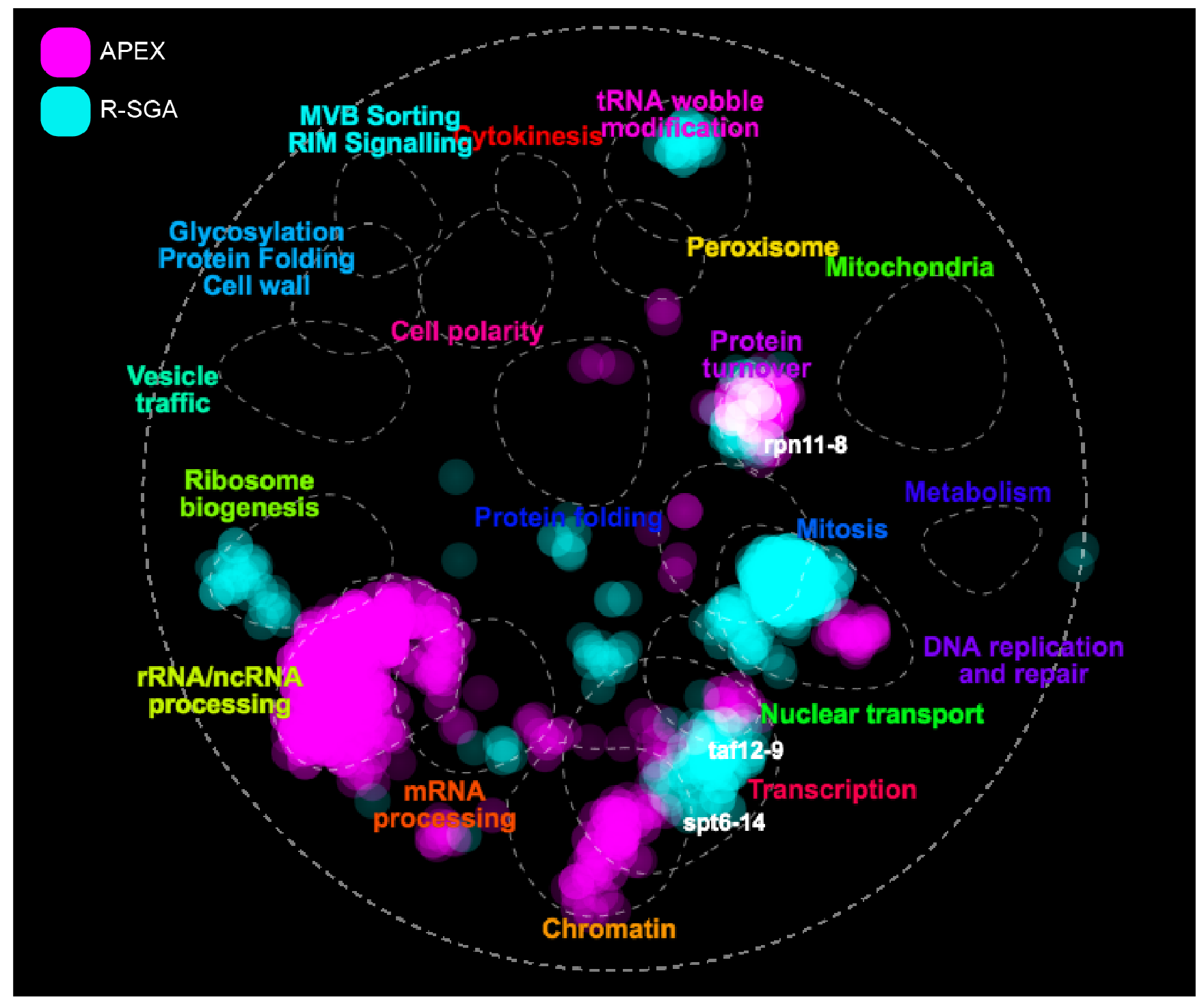
Figure S7. A global genetic interaction profile similarity network produced by The Cell Map (TheCellMap.org). The APEX identified list of nuclear interactors and the up-regulated hits from R-SGA were mapped onto the existing network to visualize localization of hits to specific sub-networks of genes. A few example genes are highlighted (spt6-14, taf12-9, rpn11-8) to demonstrate their location within interesting hubs.

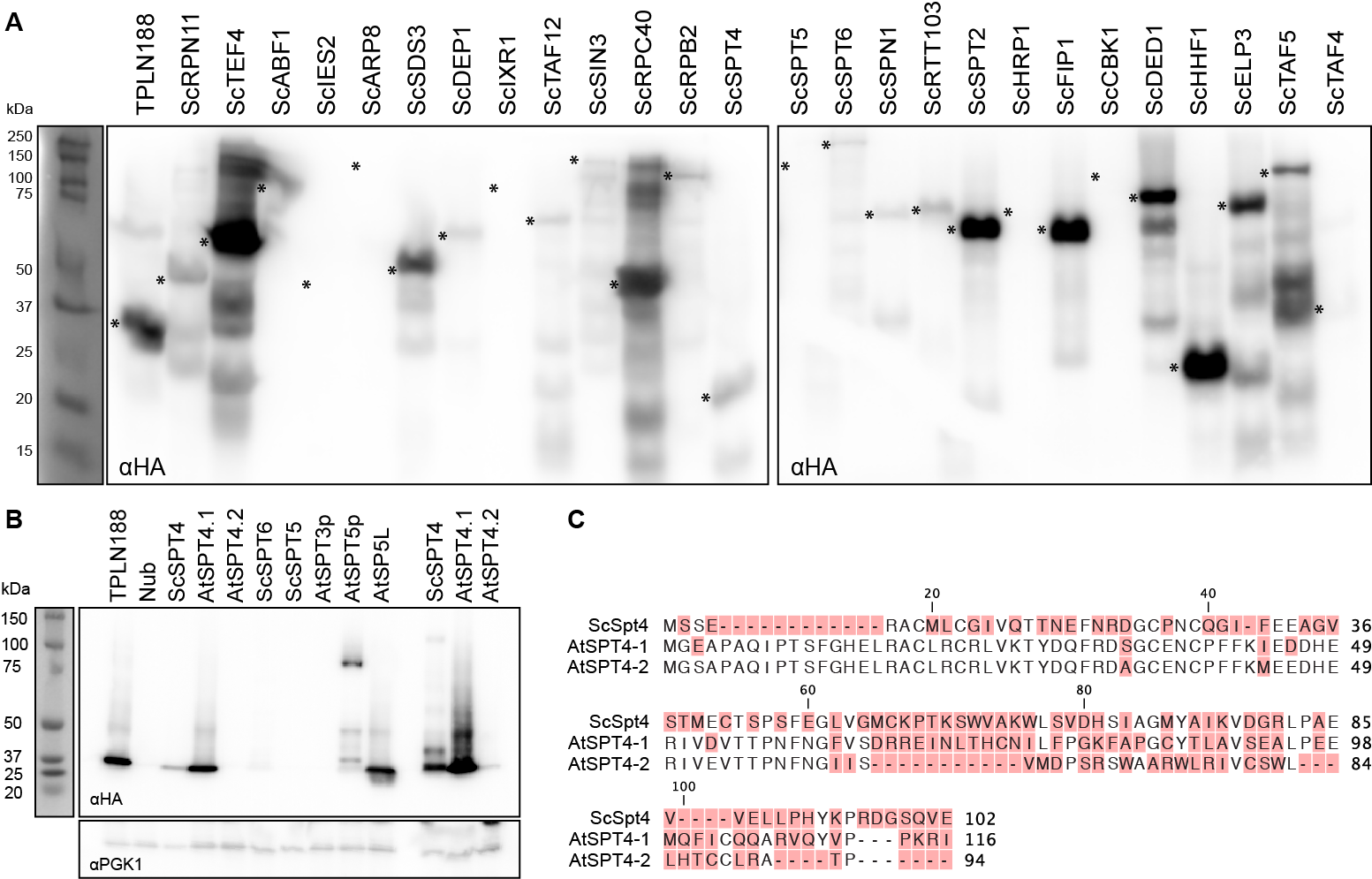

**Figure S8. TPL N-terminus interacts with ScSpt4 and AtSpt4.1 A.** Protein expression analysis by Western Blot for all tested cytoSUS targets from Figure 3D. Asterisks indicate the expected band size for the protein and are aligned to the left of the band position for each lane. **B.** Protein expression analysis by Western Blot for SPT protein expression. ScSp4, AtSPT4.1 and AtSPT4.2 were run with higher volumes of protein on the far right, demonstrating lower, but still detectable protein expression levels. **C.** Alignments of the Saccharomyces (Sc) *Arabidopsis* (At) SPT4 proteins are shown above. Non-conserved amino acids are highlighted in red.

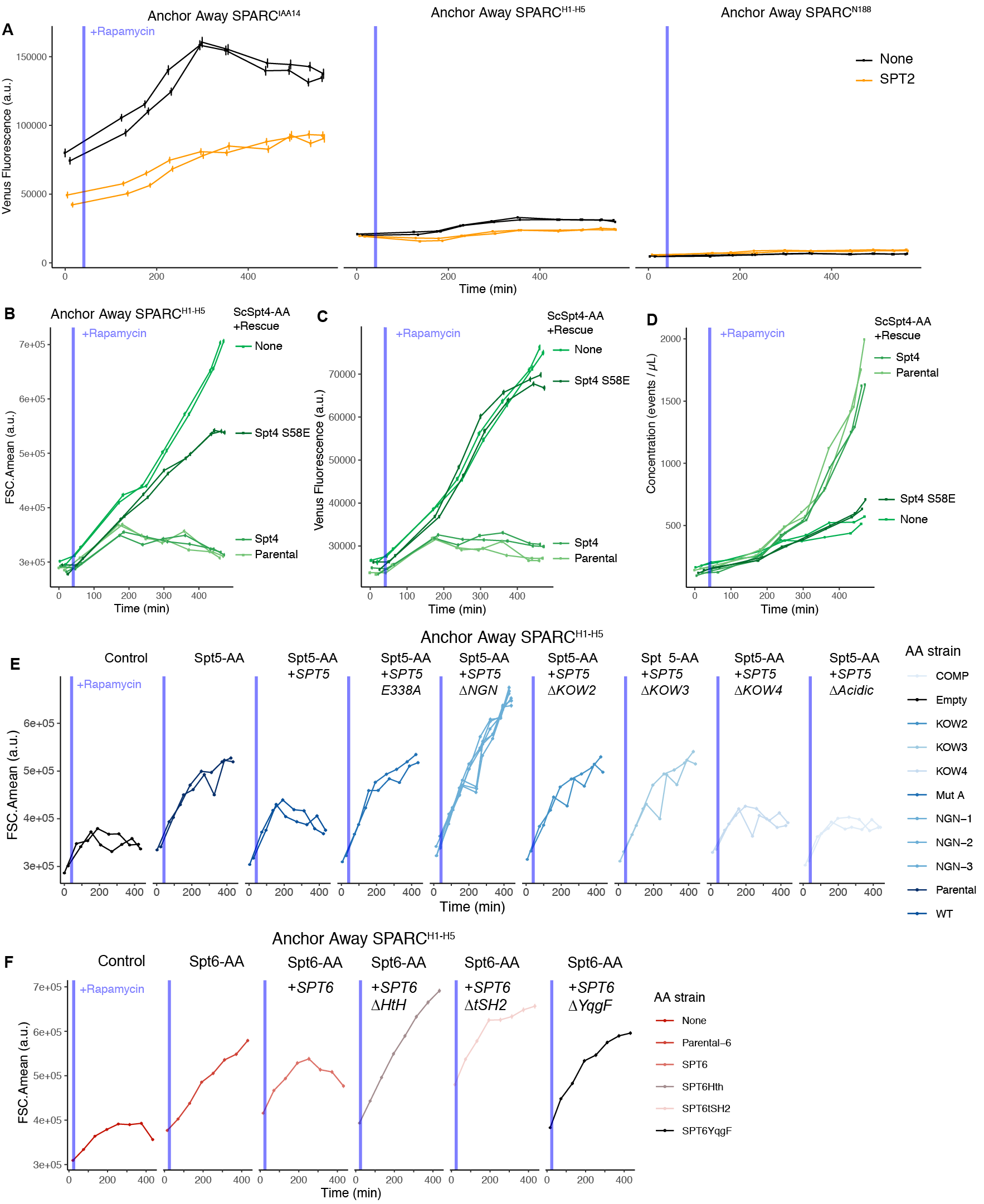

**Figure S9.** **The SPT4/SPT5/SPT6 is functionally required by TPL for repression.** **A.** Time-course flow cytometry analysis of SPARC transcription in ScSpt2 Anchor Away strains. **B-D.** Time-course flow cytometry analysis of SPARC^H1-H5^ in ScSpt4 Anchor Away strains with selected genome-integrated *ScSPT4* rescue constructs. **B**. Cell size is plotted on the y-axis as the mean forward scatter (FSCA.mean) over time on the x-axis. This demonstrates that ScSpt4 S58E has an intermediate effect on cell size compared to the ScSpt4-AA alone. **C.** Raw Venus fluorescence is plotted on the y-axis over time on the x-axis. This demonstrates that the Spt4 S58E breaks repression. **D**. Culture concentration is plotted on the y-axis over time on the x-axis. This demonstrates that ScSpt4 S58E fails to rescue the Spt4 cell division phenotype. **E**. Time-course flow cytometry analysis of SPARC^H1-H5^ in SPT5 Anchor Away strains with selected genome-integrated *ScSPT5* rescue constructs. Cell size is plotted on the y-axis as the mean forward scatter (FSCA.mean) over time on the x-axis. This demonstrates that ScSpt5 has an intermediate effect on cell size compared to the Spt4-AA alone. **F**. Time-course flow cytometry analysis of SPARC^H1-H5^ in SPT6 Anchor Away strains with selected genome-integrated *ScSPT6* rescue constructs. Cell size is plotted on the y-axis as the mean forward scatter (FSCA.mean) over time on the x-axis.

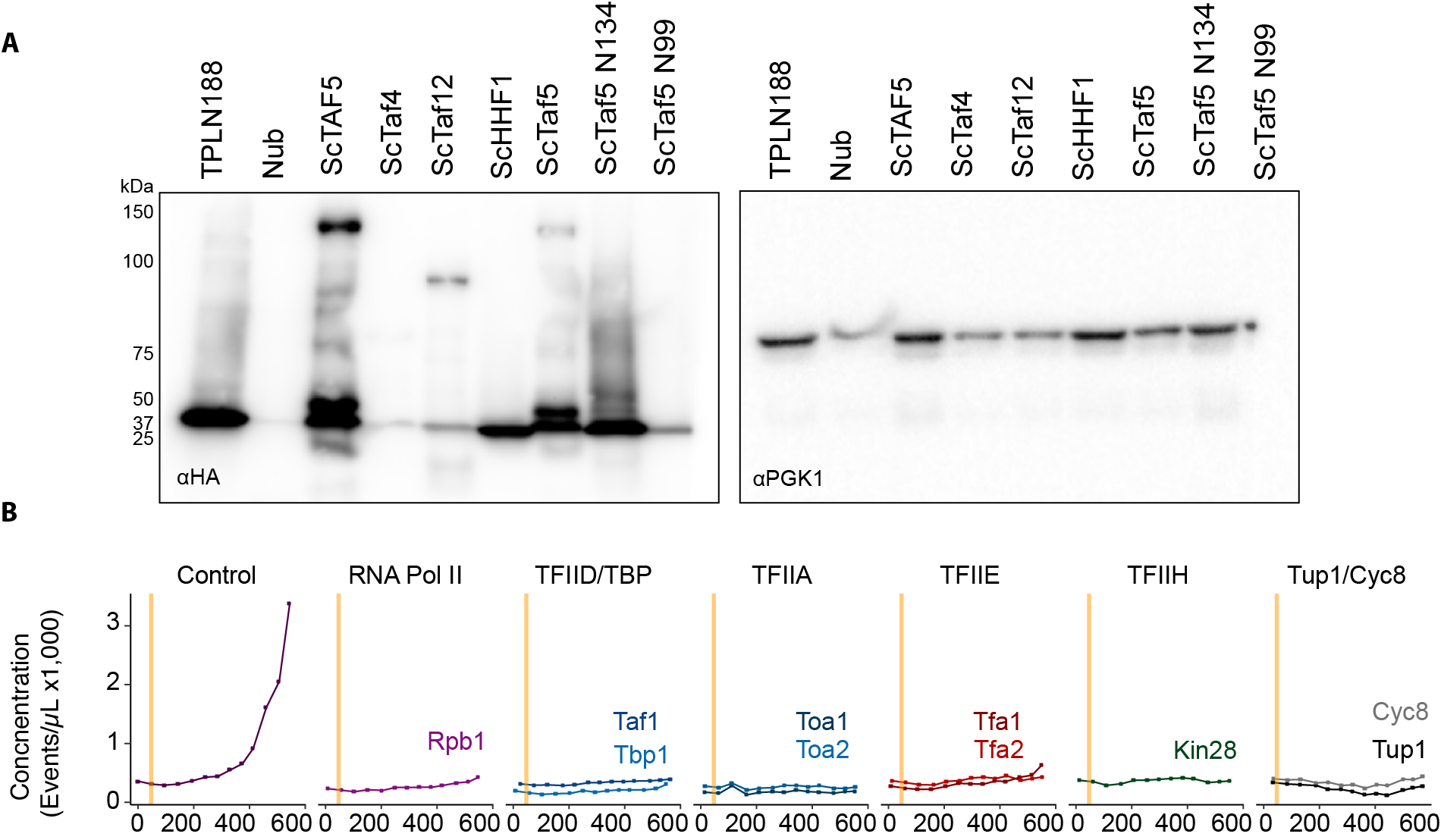

**Figure S10. A.** Protein expression analysis by Western Blot for ScTaf protein expression. **B.** Time-course flow cytometry analysis of SPARC^H1-H5^ in selected Anchor Away strains that include selected GTF components^58^. Culture concentration is plotted on the y-axis over time on the x-axis to highlight how most essential gene anchor away experiments are stymied by impacts on cell growth and division.

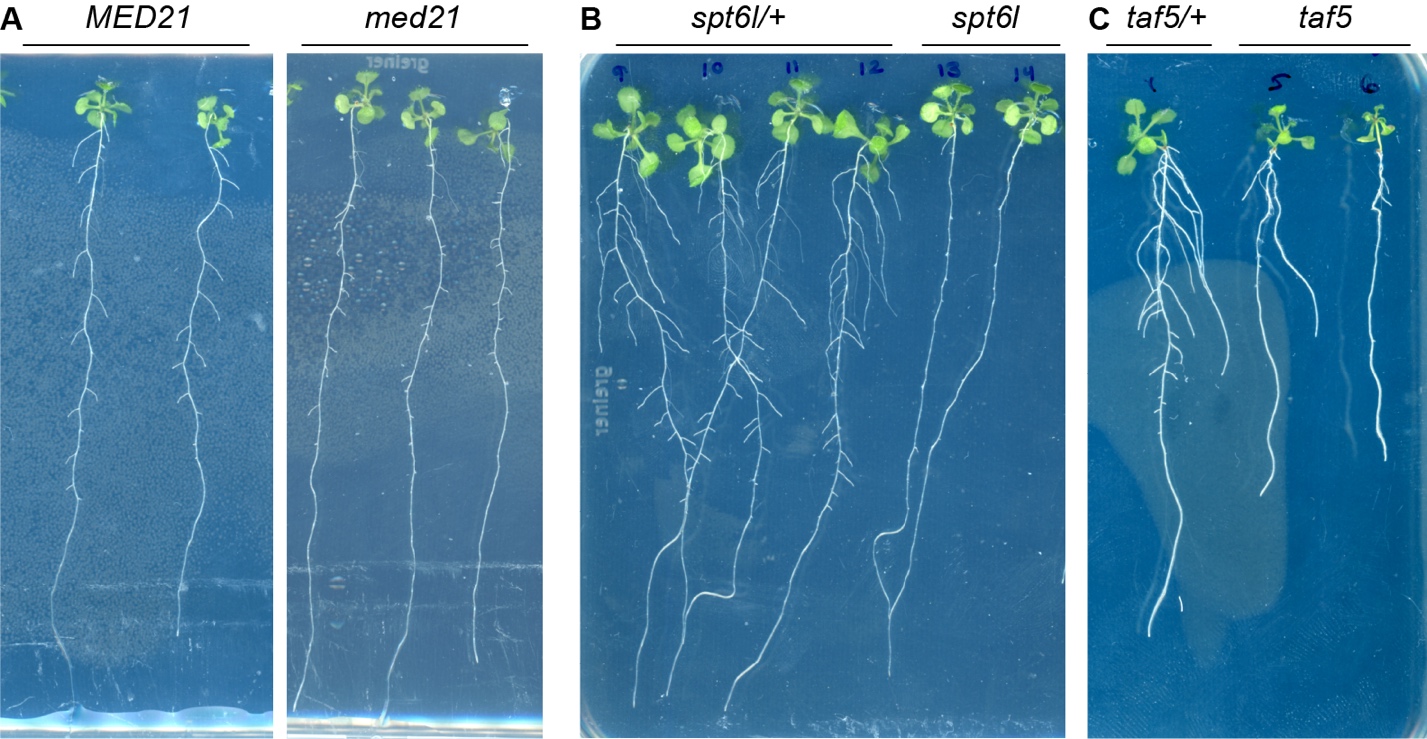

**Figure S11. Integrase switched essential lines demonstrate developmental root phenotypes.** Scanned images of seedlings grown on plates for 14 days. **A.** MED21 integrase switch lines exhibit short stubby roots that grow out from the primary root. **B.** SPT6L integrase switch lines exhibit very short roots that do not exit the primary root and are not visible in plate scans. **C.** TAF5 integrase switch lines frequently exhibit short roots that do not exit the primary root and are less visible in plate scans.
